# Evolution is coupled with branching across many granularities of life

**DOI:** 10.1101/2024.09.08.611933

**Authors:** Jordan Douglas, Remco Bouckaert, Simon C. Harris, Charles W. Carter, Peter R. Wills

**Affiliations:** Department of Physics, The University of Auckland, New Zealand; Centre for Computational Evolution, The University of Auckland, New Zealand; School of Computer Science, The University of Auckland, New Zealand; Department of Statistics, The University of Auckland, New Zealand; Department of Biochemistry and Biophysics, University of North Carolina at Chapel Hill, USA; Integrative Transcriptomics, Interfaculty Institute for Bioinformatics and Medical Informatics (IBMI), University of Tübingen, Germany

## Abstract

Across many different scales of life, the rate of evolutionary change is often accelerated at the time when one lineage splits into two. The emergence of novel protein function can be facilitated by gene duplication (neofunctionalisation); rapid morphological change is often accompanied with speciation (punctuated equilibrium); and the establishment of cultural identity is frequently driven by sociopolitical division (schismogenesis). In each case, the change resists re-homogenisation; promoting assortment into distinct lineages that are susceptible to different selective pressures, leading to rapid divergence. The traditional gradualistic view of evolution struggles to detect this phenomenon. We have devised a probabilistic framework that constructs phylogenies, tests hypotheses, and improves divergence time estimation when evolutionary bursts are present. As well as assigning a clock rate of gradual evolution to each branch of a tree, this model also assigns a spike of abrupt change, and independently estimates the contributions arising from each process. We provide evidence of abrupt evolution at the time of branching for proteins (aminoacyl-tRNA synthetases), animal morphologies (cephalopods), and human languages (Indo-European). These three cases provide unique insights: for aminoacyl-tRNA synthetases, the trees are substantially different from those obtained under gradualist models; Cephalopod morphologies are found to evolve almost exclusively through abrupt shifts; and Indo-European dispersal is estimated to have started around 6000 BCE, corroborating the recently proposed hybrid explanation. This work demonstrates a robust means for detecting burst-like processes, and advances our understanding of the link between evolutionary change and branching events. Our open-source code is available under a GPL license.

## Introduction

> “<u>There are decades where nothing happens; and there are weeks where decades happen.</u>” – Lenin.

Understanding the evolutionary relationships within groups of genes, species, or cultures has traditionally hinged upon a fundamental assumption: that evolutionary change occurs independently of branching. Under the molecular clock hypothesis, evolution occurs in a clock-like manner, ticking through time at a more-or-less constant rate.^1–3^ This view of evolution is embedded in phylogenetic clock models, including strict, relaxed, and local^4–7^ – each stemming from an assumption of extended, temporally regular, clock-like evolution; or gradualism. This view cannot detect evidence for the scenario where evolutionary change is abruptly accelerated when one lineage splits into two, or when the change is what causes the split. Rather, a phylogenetic tree is seen as a Jackson Pollock canvas that mutations are “thrown” onto, in a manner that does not perturb the tree or alter the course of evolution. Application of an inappropriate phylogenetic model is often met with unwanted consequences; divergence time estimates can become biased, and unwarranted confidence can be placed in incorrect relationships.^8–10^

In the limiting case, genetic mutations disproportionately arise in temporally abrupt bursts at the time of branching (i.e., copy errors during replication).^11^ These effects are averaged out by the molecular clock over evolutionary timescales, as shown by statistical testing.^12^ However, evolution is known often to occur in saltational leaps^13^ that are frequently, but not necessarily, tied to branching (Fig. 1). In this scenario, the gradualistic view is dubious.

**Fig. 1:**
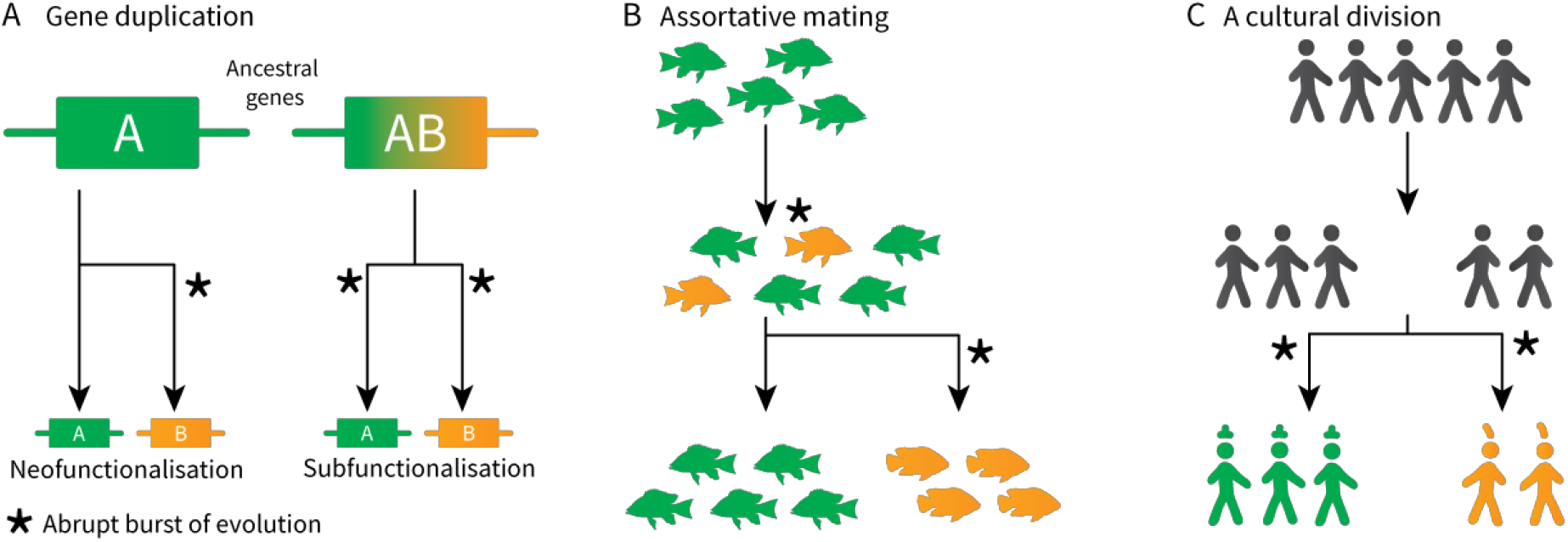
Evolutionary changes are often the driving force behind branching, or are accelerated by branching. The scenarios above depict examples of this process at molecular (A), morphological (B), and human cultural (C) scales. The coupling between evolutionary change and lineage splitting cannot be explained by clock-like evolution alone.

At the molecular level, gene duplication and lateral transfer can induce the *de novo* emergence of protein functions. In the 1930s, Haldane hypothesised that one copy of the gene will inevitably become silenced through random mutation.^14^ This means that, conditional on both copies surviving, one paralog (or both) is likely to have gained a novel selective advantage relative to the other. In some cases, one daughter copy evolves a novel function while the other retains its ancestral function (*neo-functionalisation*;^15^ Fig. 1A). In other cases, either daughter takes on a subset of the parental functions (*subfunctionalisation*^16, 17^*). Such mechanisms have given rise to a wide diversity of proteins, as exemplified by the spectrum of aminoacyl-tRNA synthetases*.^*18*^ *Environmental change might spur on further rapid adaptation; for example, antibiotic resistance*.^*19*^

*At the macroevolutionary level, Eldredge and Gould 1972 described a process they called punctuated equilibrium*,^20, 21^ whereby species undergo long periods of stasis characterised by gradual evolution, followed by short periods of abrupt evolution that are often linked to speciation. Although the theory of punctuated equilibrium was initially met with much skepticism and debate,^22–24^ it has since been quantitatively established in many systems on both genetic^25–27^ and morphological^28, 29^ scales, and rejected in others.^30, 31^ In some cases, branching accelerates the rate of species evolution, such as when a population migrates to a new habitat (adaptive radiation). In other cases, a sudden circumstantial change is what induces the branching, such as when individuals select mates with similar phenotypes (assortative mating;^32^ Fig. 1B).

At the anthropological level, human cultures may change quite abruptly when new communities are founded. This creation of a social division has been termed *schismogenesis* by Bateman 1935,^33^ and *esoterogeny* by Thurston 1987.^34^ When considering language evolution, the effect can manifest as rapid gains or losses of the *cognates* of a language (two words are considered cognate if they share common ancestry – for example “criatura” in Spanish is cognate with “creature” in English, but not “animal”). The notion of languages evolving like species – in punctuated bursts – has long been proposed as an important process in their development,^34–37^ and quantitatively demonstrated in several language families.^38, 39^ This might result from communities seeking to establish cultural identities (Fig. 1C), or as a founder effect – potentially in a new environment.

Saltation events have a structural similarity across these three cases. This similarity comes in three parts: a random heritable change, a foothold, and a fixative event that drives population splitting. First, the change appears (e.g., mutation). Second, the environmental context, either internal or external to the system, provides a foothold allowing the change to produce a newly viable variant that persists through time and avoids re-homogenisation. Third, this foothold permits a fixative process in which the change resists re-homogenisation (and entrenches heterogeneity), and the assortment persists. The differential across the two lineages means they are susceptible to novel, or previously non-existent, unique selective pressures, which will allow further random changes to accumulate and giving the appearance of an accelerated clock rate; an *epistatic ratchet*.^40^ The same pattern is seen in established models of the origin of life^41^ and genetic coding.^42^ Each is based on the splitting of initially homogeneous populations of either protocells or sets of proteins, respectively, initiated by the growth of a fluctuation (random change) away from a homogeneous ground state, through a dynamic instability where the bifurcation establishes a foothold, leading to its fixation in a new heterogeneous state.

These cases highlight the inability of gradualism to explain important evolutionary processes. Phylogenetic clock models – such as strict,^1^ relaxed,^4,5,43^ and local^5–7, 44^ – describe ways in which change occurs on average through time (Fig. 2). Strict clocks impose a fixed rate of change, while relaxed (uncorrelated) clock rates vary independently across lineages. “Relaxed clock” is usually shorthand for “relaxed gradual clock”, but abrupt changes at branch points might also be strict (with each branch experiencing an instantaneous “spike” of constant magnitude^25, 39^) or relaxed (with each branch having a spike of independent magnitude^27^).

**Fig. 2:**
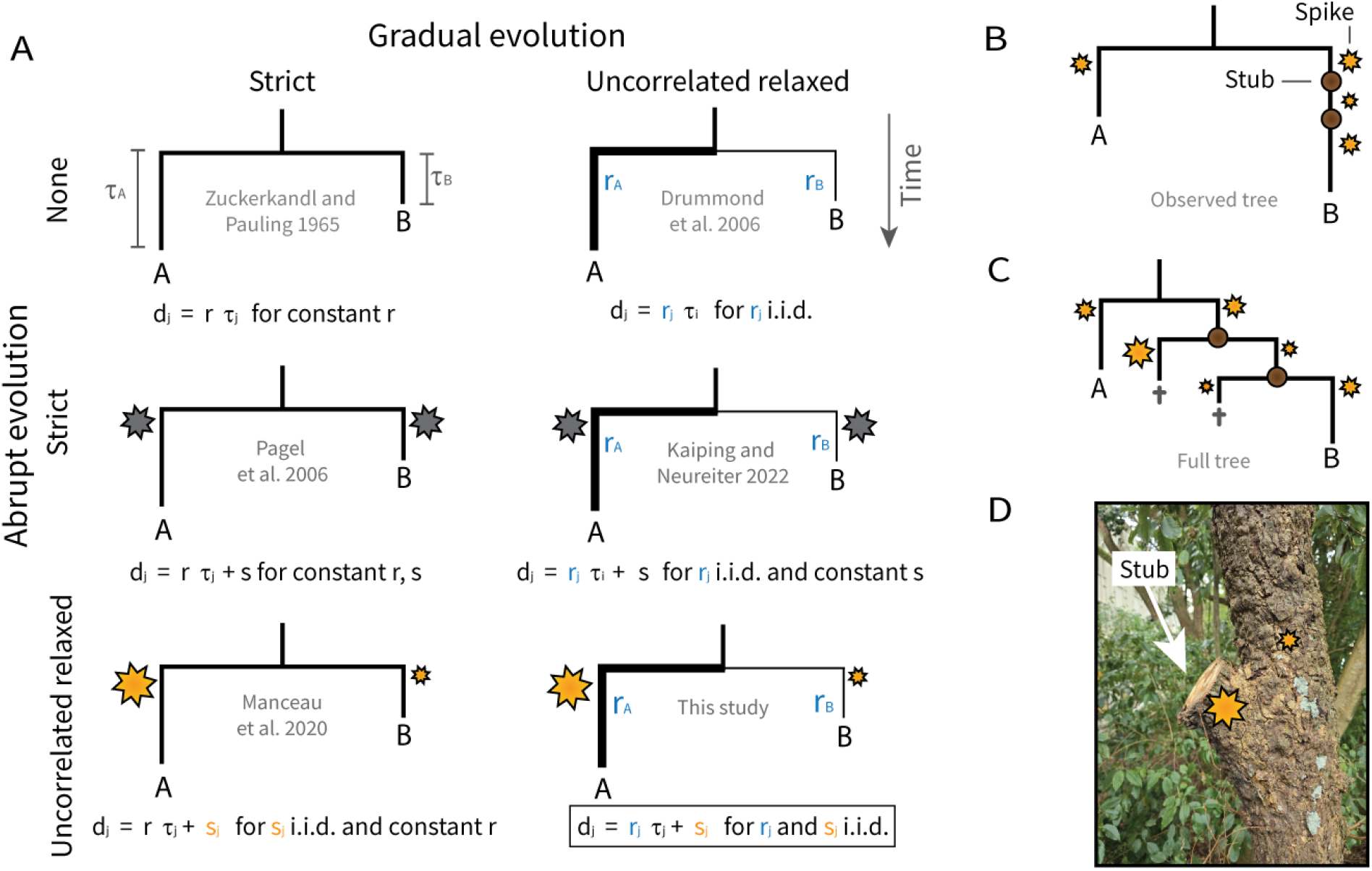
Gradual and abrupt models of evolution. A: In the (uncorrelated) relaxed models of gradual evolution, relative branch rates (line widths) are *a priori* assumed to be independent and identically distributed (i.i.d.). The relaxed models of abrupt evolution assume that each branch is characterised by an instantaneous spike of evolution (stars), and the sizes of those spikes are also i.i.d. Note that Manceau et al. 2020^27^ assumed that branch spikes take a more restricted range of values, rather than the continuous magnitudes depicted here. Correlated or local clock models represent a midpoint between strict and uncorrelated. B-C: If one assumes that each branch is associated with an evolutionary spike, then unobserved speciation events (i.e., stubs) must be accounted for. These unsampled taxa may leave behind a footprint on those that were sampled, by inducing hidden evolutionary bursts. D: A stub on a wooden tree at the University of Auckland.

To investigate the link between branching and change, we provide a probabilistic method for building phylogenies and testing abrupt evolution. Expanding on existing methods,^25, 27, 39, 45^ our model captures both abrupt (relaxed) and gradual (relaxed) evolution. If branching and evolution are indeed tightly coupled, then unobserved speciation events may have left behind a footprint on the lineages that have been observed.^27, 39, 45^ We assume that each lineage experiences a rapid evolutionary spike, whose size is informed by the number of unobserved bifurcations along that lineage (i.e., the number of *stubs*; Fig. 2). Despite the model’s high dimensionality, it has many attractive properties, such as statistical consistency, low covariation between rates and spikes, and its ability to recover the known model from simulated data. We hypothesise that abrupt evolution can occur anywhere in nature; demonstrating the process in genes, morphologies, and languages.

## Results

In the following sections, we demonstrate what is gained by considering the effects of abrupt evolution in addition to gradual change. We do this by benchmarking the proposed “gradual+abrupt” clock model (with both components relaxed) against a standard “gradual” relaxed clock model.^4,43^ Gradualistic divergence can be thought of as independent diffusion, and abrupt change as a transient repulsion that drives the lineages apart even further. The alternatives are compared using Bayesian model averaging.^46^ Much like the Akaike and Bayesian information criteria, Bayesian model averaging penalises overparameterised models, and selects against an overly complex model when it has little to offer. As shown in Fig. 2A (bottom right), the new model does have a large parameter space, with each branch consisting of a length (time), a rate (gradual evolution), and a spike (abrupt evolution). To further dampen the effect of overparameterisation, these three terms are constrained within a narrow range of values, by assigning them prior distributions with low variances. Tree branching follows a fossilised-birth-death process,^47^ with the number of stubs per-lineage also being estimated. We first validate and characterise this method through simulation studies, and then investigate fourteen empirical datasets, three of which are covered in-depth.

### Simulation studies

The proposed method was implemented in a probabilistic setting. In this way, parameters, trees, and sequences can be directly simulated under a model the same, or different from, that used during inference. We used this property to validate our new method through coverage simulation studies.^48^ This was achieved by i) sampling parameters and trees *θ* from a probability distribution; ii) simulating a multiple sequence alignment *D* under *θ*; iii) performing Markov chain Monte Carlo (MCMC) on data *D* to obtain parameter estimates 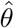; and lastly iv) comparing *θ* with estimates 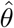. These four steps were repeated multiple times. When the model is correctly implemented, one would expect the true value of a parameter to reside in its 95% credible interval in approximately 95% of all replicates (in which case we say it has 95% coverage). These experiments confirmed the usefulness and correctness of the method, whose estimates were well-correlated with the true values and had around 95% coverage (Supporting Information and Fig. 3C).

**Fig. 3:**
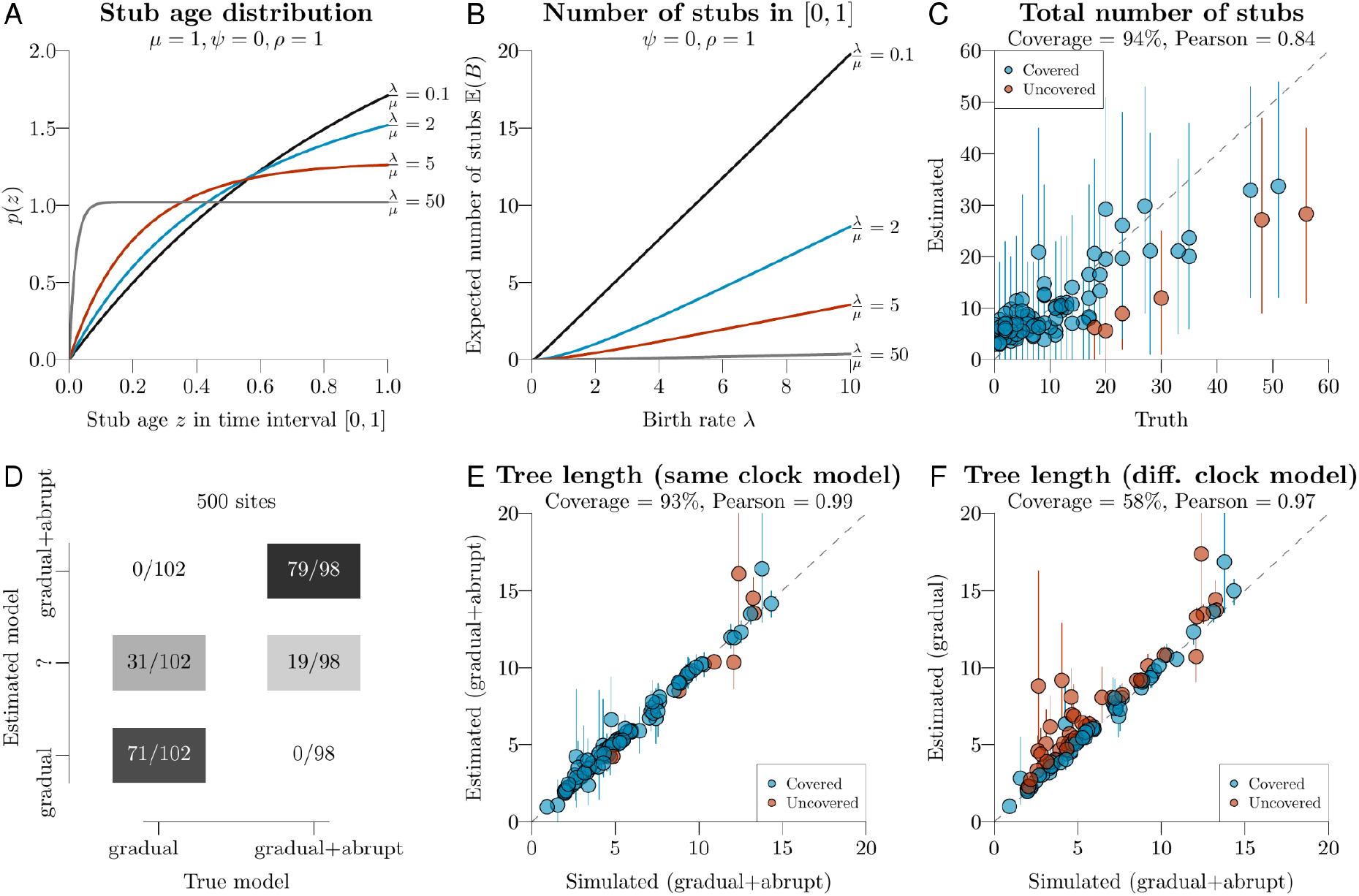
Characterising the probabilistic properties of the model. A: Most stubs occur near the start of a lineage. B: The number of stubs along a lineage depends on branching parameters (birth rate *λ*, death rate *µ*, historical sampling rate *ψ*, and contemporaneous sampling probability *ρ*). C: Coverage simulation study for estimating the number of stubs. Each point is one MCMC chain and one simulated set of parameters; mean estimates are depicted by circles, and 95% credible intervals by vertical lines. The trees were fixed at their true values, while branching parameters and number of stubsperlineage were estimated during MCMC. D: Bayesian model averaging - with 500 sites and a mean spike size of *S*_*µ*_ = 0.01, the true clock model can be recovered from simulated data; the middle row ‘?’ denotes the ambiguous case when the inferred model has a Bayes factor of less than 10 (i.e., 0.09 *< p*(𝕀_*c*_ = 1) *<* 0.91). E: Each point is a single MCMC chain running on a respective DNA dataset simulated under the gradual+abrupt clock model. The tree length is usually recovered within the 95% credible interval. F: Using the same 100 datasets as E, here MCMC was instead performed under the gradual clock model. This experiment suggests that the gradual clock model overestimates divergence times on datasets that are generated by abrupt evolution.

Second, we evaluated the method’s ability to identify the true clock model using Bayesian model averaging. To do this, we simulated data under two models (gradual vs gradual+abrupt) and estimated the clock model indicator 𝕀_*c*_ ∈ {0, 1}. A Bayes factor of 10 indicates “strong” support in favour of one hypothesis over another,^49^ corresponding to *p*(𝕀_*c*_ = 1) = 0.91 (or 0.09) when the two hypotheses are equal *a priori*. Using this threshold, and “true” spikes averaging 0.01 substitutions per site per bifurcation, the method recovered the true clock model in 150/200 simulated alignments (with 500 sites) and was uncertain in the remaining 50/200 (Fig. 3D). On smaller datasets with just 20 and 100 sites, the clock model was correctly identified 13/200 and 81/200 times respectively, showing that accuracy improves with increasing data volume. The wrong model was never selected. We also assessed the effect of model misspecification – where the prior distributions used during inference differed vastly from those used during simulation. Even under these conditions, the clock model was still rarely wrong, and support in favour of the correct model grew with increasing alignment length (Fig. S4). These results are reassuring; they confirm the method i) is statistically consistent, ii) is reliable at hypothesis testing even when there is no prior information, and iii) does not have a proclivity to “overfit” using the extra parameters of the larger model.

Lastly, we assessed the effect of model misspecification on divergence time estimation (Fig. 3E-F). To do this, we estimated divergence times for each clock model on datasets that were simulated under the gradual+abrupt model. The 100 trees used here were again seriallysampled timetrees. The sequence alignments (with 100 sites) were simulated with an average spike size of 0.02 substitutions per site per bifurcation (abrupt evolution), and an average clock rate of 1 substitution per site per unit of time (gradual evolution). We then compared the known tree length (i.e., the sum of all branch lengths) with the tree length inferred under either clock model (gradual vs gradual+abrupt). Unsurprisingly, the gradual+abrupt model was able to infer the correct tree length (Fig. 3E), whereas, branch lengths (time) were overestimated (by 16%) under the gradual model (Fig. 3F).

### Empirical datasets

We assessed these two views of evolution on fourteen empirical datasets. First, we screened the method on a widely used collection of eleven nucleotide datasets DS1-11.^50^ Using a Bayes factor threshold of 10, five of these datasets favoured the gradual+abrupt model,^51–55^ three rejected it,^56–58^ and the remainder were less certain.^59–61^ The datasets that did support the new model yielded mean spike sizes *S*_*µ*_ ranging from 0.005 to 0.02, while those that rejected this model yielded smaller estimates (Table S1).

Second, we took a deeper dive into three empirical case studies: aminoacyl-tRNA synthetases (protein sequences), cephalopods (animal morphologies), and Indo-European languages (cognates). The gradual+abrupt evolution model was overwhelmingly favoured, with estimated probability *p*(𝕀_*c*_ = 1) = 1 in each case, suggesting that these systems underwent significant rapid changes. The trees in Fig. 4 summarise the gradual+abrupt tree posterior distributions shown in Fig. 5, which reveal that the competing models yielded divergent clade probabilities, to statistically significant and oftentimes substantial levels. This effect was strongest for the protein dataset. As shown in Table 1, the average spike size varied considerably between the three datasets, from 0.001 substitutions per site per bifurcation for the language dataset, to 0.065 for the proteins. The non-zero spike sizes resulted in less change being explained by gradual evolution. Therefore the estimated gradual clock rates were lower (with the exception of the protein dataset, whose clock rate was fixed due to the absence of temporal calibration). The data provided no signal for estimating the shape of the spike Gamma distribution *S*_*α*_. In the most extreme case, the clock rate was two orders of magnitude smaller when abrupt evolution was present (cephalopods).

**Table 1:** Comparison of the clock models, gradual (G) and gradual+abrupt (G+A), on three empirical datasets.

| Dataset | Clock model | Tree height | Tree length | Clock rate<br>$\mu_c$ | Spike mean<br>$S_\mu$ | Spike shape<br>$S_\alpha$ | Stubs | $\rho_{sr}$ |
| --- | --- | --- | --- | --- | --- | --- | --- | --- |
| aaRS | G | 3.98<br>(3.83, 4.13) | 109<br>(99.4, 120) | 1 | - | - | 16<br>[2, 36] | - |
|  | G+A | 3.96<br>(3.8, 4.1) | 89.5<br>(77.5, 102) | 1 | 0.065<br>(0.036, 0.094) | 2.2<br>(0.82, 4) | 12.4<br>[1, 27] | -0.12 |
| Cephalopods | G | 494<br>(469, 553) | 3800<br>(3230, 4400) | 0.0044<br>(0.0022, 0.0071) | - | - | 186<br>[85, 296] | - |
|  | G+A | 500<br>(469, 567) | 3680<br>(3070, 4320) | 2.2e-05<br>(3.2e-09, 9.3e-05) | 0.045<br>(0.025, 0.068) | 2.3<br>(0.96, 4) | 111<br>[60, 163] | -0.00081 |
| Indo-European | G | 7.62<br>(5.99, 9.36) | 186<br>(164, 207) | 0.011<br>(0.0066, 0.017) | - | - | 188<br>[138, 241] | - |
|  | G+A | 7.92<br>(6.19, 10) | 184<br>(159, 208) | 0.0094<br>(0.0036, 0.019) | 0.0014<br>(0.00054, 0.0031) | 1.8<br>(0.61, 3.1) | 192<br>[139, 248] | -0.17 |

Parameter mean estimates (and 95% credible intervals) are shown. Tree heights and lengths are in units of substitutions per site (aaRS), Ma (cephalopods), and Ka (Indo-European). Therefore, clock rates are, respectively, in units of amino acid substitutions per site, trait changes per million years, and cognate changes per thousand years. $\rho_{sr}$ is the average Pearson correlation between the estimated spike and rate of each branch.

**Fig. 4:**
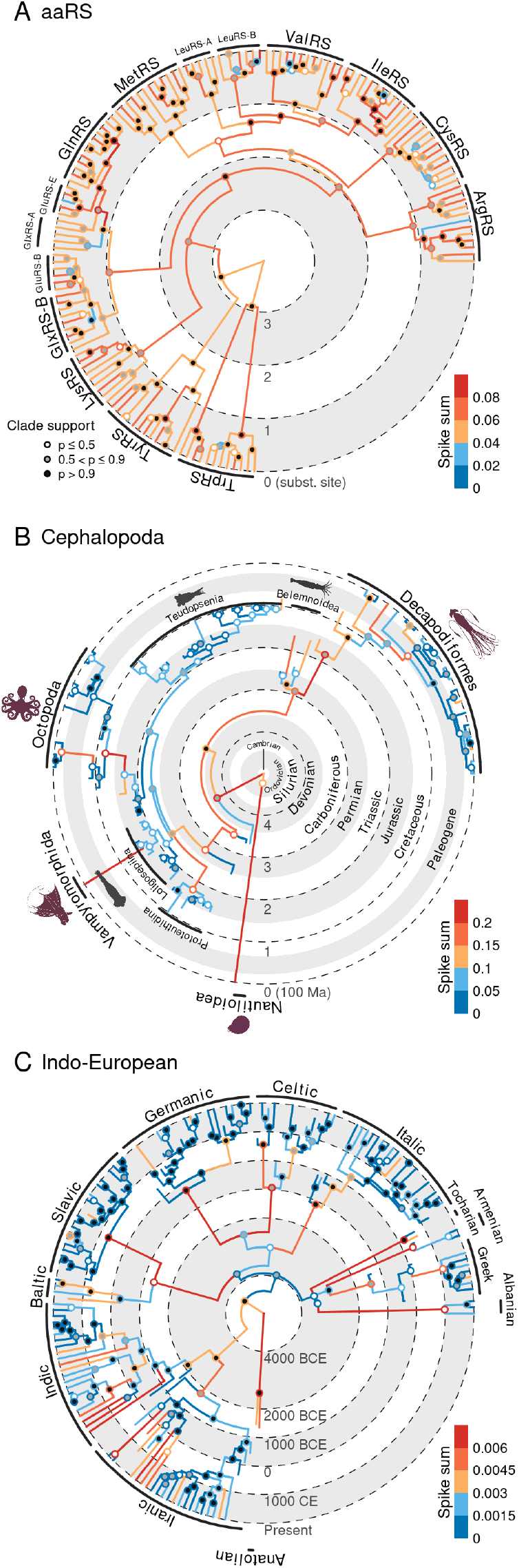
Phylogenies inferred under the gradual+abrupt clock model. The three point estimates summarise their respective tree posterior distributions using the CCD0 method.^75^ Median estimates of branch spike sums are shown.

**Fig. 5:**
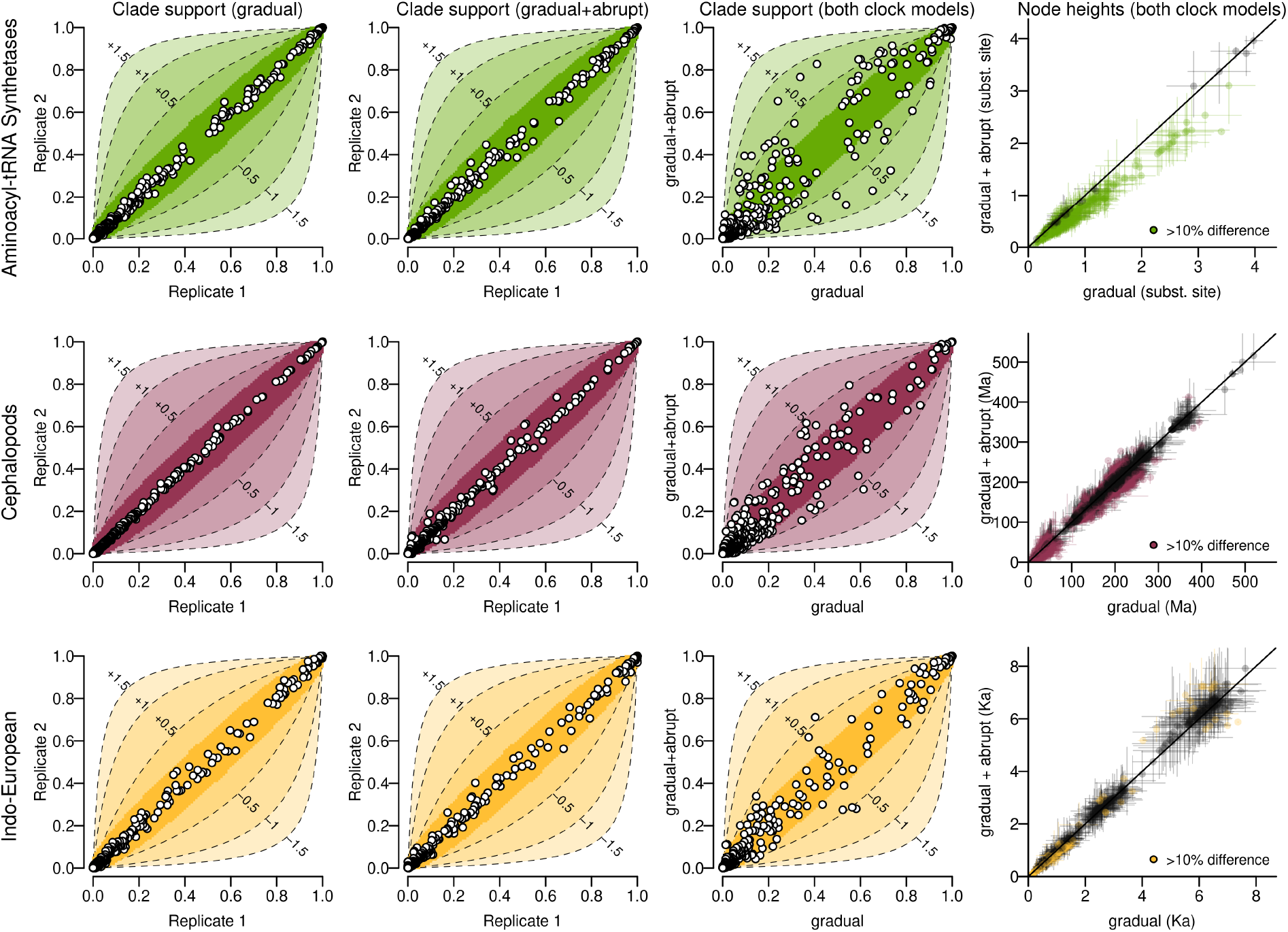
Comparison of tree posterior distributions across two clock models. Each point is a clade found in the posterior distribution of one or both MCMC chains. Clade supports are shown on cocoa-fruit plots, on columns 1–3. The first two columns depict two independent MCMC replicates, whose clade supports should lie close to the diagonal if the two chains have indeed converged to the same distribution, shown by statistically-insignificant zone (inner solid-line contour). The third column compares both clock models. Many clades are associated with substantial changes in log Bayes factor between the two models (outer dashed-line contours). Please refer to Materials and Methods for further details. Column 4 compares node heights estimates across both modes; points are means and lines are 95% credible intervals.

There is clearly a degree of covariation in estimating rates and spikes simultaneously on each branch. However, as Table 1 and Table S1 both show, the (negative) correlations between the two estimates are quite modest. This further corroborates the notion that rates and spikes are capturing distinguishable processes; justifying the additional parameters.

### Protein study: Aminoacyl-tRNA synthetases

The aminoacyl-tRNA synthetases (aaRS) are a large group of enzymes that implement the genetic code by attaching amino acids to their cognate tRNAs.^62^ The complete set of aaRS families, which enable the coding of twenty-plus canonical amino acids, emerged from a series of ancient structural and functional diversification events that can be traced back to the earliest stages of life on Earth before the last universal common ancestor.^18, 63, 64^ Therefore, it is unsurprising to find that so much of their evolution has occurred in sudden bursts, previously described as quasi-species bifurcations.^65^

We considered a previously published curated alignment of aaRS three-dimensional structural models,^66^ consisting of 494 amino acid sites from 142 Class I aaRS catalytic domains sourced from bacteria, archaea, eukaryotes, and viruses. The two clock models produced substantially different clade posterior supports (Fig. 5), which were not only statistically significant, but also induced large changes to clade Bayes factors. As shown in Table 1, these proteins have undergone considerable mutational saturation over their presumed 4 billion years of evolution, with an estimated tree height of around 4 substitutions per site. The gradual model overestimated the tree length by around 22%, relative to the gradual+abrupt model, reflecting the large mutational spikes, which saw one in every fourteen amino acids change at each bifurcation. As shown in our phylogenetic tree (Fig. 4), most lineages, young and old, were associated with sizeable spikes, notably those that gave rise to aaRS families; including arginyl-tRNA synthetase (ArgRS), lysyl-tRNA synthetase (LysRS), and isoleucyl-tRNA synthetase (IleRS). The relationships between the families are similar to those of our previous tree, which additionally took structural insertion modules into account.^18^ An even more reliable phylogeny may be obtained by accounting for both abrupt evolution and the gain of insertion modules. It is noted that, while the abrupt model does indeed give a better fit than the standard gradual model, it still makes the critical assumption of the amino acid alphabet being fixed to twenty characters through time; an assumption violated by the translational reflexivity unique to aaRS synthesis.^65^

### Morphology study: Cephalopods

The Cephalopoda are a class of molluscs characterised by their soft bodies and variable numbers of arms and tentacles.^67^ The ingroup (Coleoidea) showcases advanced intelligence and mimicry skills,^68, 69^ and includes the tenarmed Decapodiformes (squids and cuttlefish), and the eightarmed Octapoda (octo-puses) and Vampyromorphida (vampire squid; *Vampyroteuthis infernalis*). The outgroup (Nautiloidea) consists of a primitive group of molluscs, often regarded as “living fossils”,^70^ with a coiled shell and up to a few dozen appendages.

Whalen and Landman 2022^71^ compiled a dataset of cephalopod morphological traits, detailing both living organisms and fossils. These 153 discrete traits describe shell shapes, tentacle structures, the number of fins, and various other properties of 27 living cephalopods and 52 fossils. In contrast to the aaRS case, the data characterising these taxa contains temporal information; the oldest taxon being afossilised *Nautilus Pompilius* specimen dated at 469 million years ago. The authors used this dataset to build a phylogenetic tree under a fossilised birth-death tree prior and a strict (gradual) clock model.

Using the same dataset, we found that gradual evolutionary clock rate estimates varied by two orders of magnitude between the two models (Table 1). The gradual+abrupt model suggests that gradual evolution played a trivial role in producing the morphologies of these taxa (1.7 of 153 traits are estimated to change over 500 million years), which predominantly changed in rapid bursts (an average of 6.9 of 153 changes per speciation event). Hence, the “gradual+abrupt” model is effectively more of an “abrupt” model in this case. As a result, the spikes and rates on each branch were uncorrelated (*ρ*_*sr*_ ≈ 0). The two clock models also yielded different clade probabilities, at a statistically significant level, but the differences are not as extreme as for the aaRS (Fig. 5). As shown in Fig. 4, there are two lineages associated with disproportionately large levels of abrupt change: the lineages that gave rise to i) the coleoid ingroup during the late Cambrian period [average spike sum: 0.37 (0.08, 0.76)], and ii) the vampire squid during the Jurassic period [0.31 (0.13, 0.57)]. We note that the large spike sum on the nautoloid outgroup lineage is a false positive reflecting the large number of stubs required to account for the putative evolutionary relationship between the outgroup and the Coleoidea ingroup; in reality this living specimen has the same morphology as its fossilised ancestor near the root of the tree (but with several “missing traits” that are ignored by the Mk Lewis model^72^). This work further corroborates the phylogeny produced by Whalen and Landman 2022,^71^ and is generally consistent with the picture derived from molecular data.^73, 74^

### Language study: Indo-European

The Indo-European languages – including English, Spanish, Hindi, and Farsi – are spoken by around half of the world’s population as a first language.^76^ The origins of this language family have been contested for over 200 years. The Steppe hypothesis proposes an expansion from the Pontic-Caspian Steppe (Eastern Europe and Central Asia) some time after 4,500 BCE.^77^ The farming hypothesis suggests that Indo-European dispersed from the Fertile Crescent (North Africa and Eurasia) around 6,500 - 7,500 BCE.^78^

Heggarty et al. 2023 compiled an extensive dataset containing 4990 cognates across 109 present-day and 52 ancient Indo-European languages.^79^ The oldest languages (such as Hittite, Luvian, and Mycenaean Greek) date to before 1000 BCE. Using a relaxed (gradual) clock model, they estimated that Indo-European dispersal started around 6,150 BCE; an age inconsistent with both the Steppe and farming hypotheses. Rather, they propose a hybrid explanation. A recent study points out that this hypothesis is difficult to reconcile with genetic data.^80^

We reinterpreted the Heggarty dataset by accounting for abrupt evolution. There was strong support for punctuated bursts of evolution, which can explain effects that gradualistic and covarion^81^ processes were unable to. As shown in Fig. 4, the majority of recorded ancient languages are comparatively recent (e.g., younger than 1,000 BCE). Consequently, the earlier lineages have more stubs (reflecting fewer samples), and therefore they conceal many hidden bursts of evolution. To reduce *a priori* analytical bias toward either the Steppe or farming hypotheses, we chose a tree prior distribution with an average height of 5,100, and a 95% credible interval of 2,760 – 7,700 BCE. The tree height was then estimated at around 5,600 (gradual) and 5,900 (gradual+abrupt) BCE (Table 1). An “epoch model”^82, 83^ would overcome the limitations imposed by our assumption, not applied in the previous study, of homogeneous branching through time.^81^ Taken together, our results further corroborate the hybrid hypothesis. No doubt their model will remain a subject of debate by linguists in the years to come.

## Discussion

The coupling between branching and saltation plays an important role in shaping the course of evolution across varying granularities of life, from genes to species to human civilisations. We hypothesise these two processes are frequently intertwined at other scales too, such as cellular, developmental, and ecological. While the underlying mechanisms will clearly differ at each scale, they all share a structural commonality, involving i) a random heritable change, that ii) survives long enough to establish a foothold, for iii) permitting a fixative event in which the change resists re-homogenisation, thereby coupling the fixation of complementary changes that promote splitting into distinct populations. Under stable conditions, there is a tendency for re-homogenisation,^84^ favouring gradualistic processes. However, in conditions of environmental or selective instability (perhaps induced by the change^40^), these steps might iteratively cycle, leading to the appearance of a transiently accelerated clock rate.

In this view of evolution, one must consider lineages that are extinct or unsampled. These unobserved bifurcation events, or stubs, left behind footprints on those that did survive.^27, 39, 45^ Such hidden speciation events, and therefore hidden bursts of abrupt evolution, are most prevalent among older lineages (Fig. 3, 4). The view of gradualism, by contrast, is generally agnostic concerning this process.^4–7^

To investigate this phenomenon, we formulated a probabilistic framework for building phylogenies and testing for abrupt evolution in a single joint analysis, without the need for marginal likelihood approximation.^85^ We demonstrated that this is a statistically consistent and reliable means of hypothesis testing, even when no prior information is available. As shown in Fig. 2, the model has a large parameter space. To remedy this concern, we i) placed strong prior constraints on branch lengths, rates, and spikes, ii) employed Bayesian model averaging to disqualify extra parameters when they are not needed, and iii) used state-of-the-art proposal kernels that traverse this multidimensional space effectively.^43, 83, 86–89^ Our experiments all indicate that these efforts were successful. Given its complex nature, the model behaves surprisingly well on both simulated and empirical datasets.

This work stands apart from existing methods. Pagel et al. 2006 provided a post-processing regression tool for testing punctuated equilibrium on a predetermined tree, which might not be built with abrupt evolution in mind.^25^ Manceau et al. 2020 produced a method that jointly infers spikes and other parameters, while testing abrupt evolution on a branch-by-branch basis, but also on a fixed tree.^27^ Most recently, Kaiping and Neureiter 2022 considered a strict-abrupt relaxed-gradual clock model that jointly infers the tree.^39^ Our approach combines and expands upon the best from these previous efforts, as it i) enables testing for abrupt evolution, ii) jointly infers the tree with the spike model, and iii) uses the most general of clock models (Fig. 2). When the abrupt evolution hypothesis is rejected, the algorithm falls back on the relaxed clock,^4,43^ but with the added advantage of estimating stubs.

Understanding evolutionary saltation can unravel powerful insights, as we demonstrated for the primordial differentiation of aminoacyl-tRNA synthetase enzymes, the diversification of cephalopods, and the dispersal of Indo-European languages. In each case, there was strong support for abrupt evolution. However, the impact was quite different in each scenario. First, this approach overcame biases when estimating divergence times and tree topologies, most notably in the aminoacyl-tRNA synthetases; likely reflecting their extensive structural and functional diversification over the past ∼ 4 Ga.^18, 63^ As outlined in Materials and Methods, an increase in probability from 0.9 to 0.99 has more value than an increase from 0.5 to 0.59; depicted by the cocoa-pod shaped contours in Fig. 5. Second, the (gradual) clock rate is slower when abrupt evolution is accounted for (Table 1), especially the cephalopod clock rate, which was estimated to be two orders of magnitude slower; suggesting their morphological evolution was almost exclusively governed by sudden bursts of adaptive radiation. Third, spike magnitudes varied considerably; with the aaRS (and cephalopods) seeing one in fifteen (twenty-two) amino acids (traits) change at each bifurcation event, but Indo-European languages just one per seven-hundred. The magnitudes of these estimates, as well as those in standard benchmark datasets (Table S1; DS1-11^50^), are in line with previous studies.^25, 27, 39^ Accounting for abrupt evolution does not always alter the overall conclusion to be drawn, in which case it establishes robustness; as it did for estimating the origin of Indo-European languages.

There exist numerous avenues for future investigation. This approach offered idiosyncratic outcomes in each case study, and may behave differently yet again in other settings; potentially illuminating new modes of saltation. This study assumed that speciation and extinction occur homogeneously through time, however an enhancement would involve varying these processes across epochs^82^ or among lineages.^90^ Future investigations might also explore the possibility of further integrating branching and evolution, for instance, by incorporating stub-awareness into relaxed or local models of gradual evolution,^4–7^ coalescent models including the multispecies coalescent,^91, 92^ or at the task of species delimitation^93^ or ancestral reconstruction.^17^ Taken together, this work places us one step further from treating evolutionary change as a paint bucket, and one step toward a sculpting chisel.

## Materials and Methods

### Probability density function of a tree and its stubs

Let 𝒯 be a binary rooted time-tree. We will assume that branching in 𝒯 follows a fossilised birth-death process, as described by Stadler 2010.^47^ In this model, a lineage will either branch into two (at rate *λ*) or become extinct (at rate *µ*). Ancestral lineages are independently sampled at rate *ψ*, while extant lineages are sampled with probability *ρ*. In contrast to previous approaches that are often employed in epidemiology,^88, 94^ here we assume that sampling an ancestral lineage does not “remove” the lineage from the tree. We will also assume sub or supercritical conditions, i.e., *λ* ≠ *µ*.

We will describe time in the reverse direction, i.e., age or height. Suppose that 𝒯contains *n* extant lineages (at age 0), *m* extinct lineages (at ages **y** *>* 0), and *k* sampled ancestral lineages. There are *n* + *m* − 1 internal nodes (with ages **x** *>* **0**), including the root at age *x*_1_. As described by Eqn (9) of Stadler 2010, the probability density of observing tree 𝒯 (conditional on *n*) is:

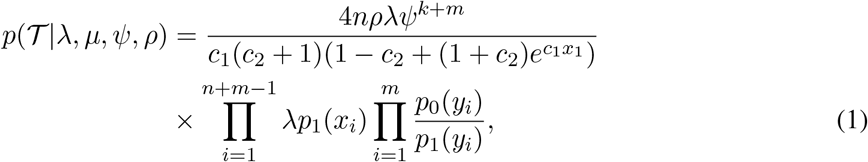

where *p*_0_(*t*) and *p*_1_(*t*) are the probabilities of an individual at age *t* respectively having 0 and 1 sampled descendants:

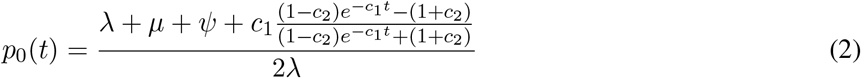

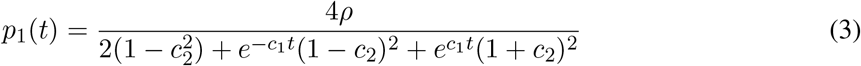

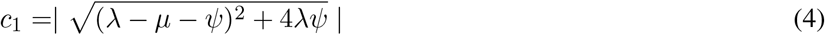

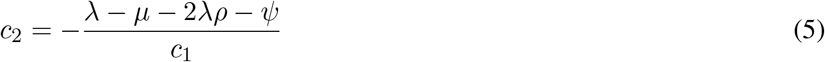

Now, let ℱ be the full phylogeny of 𝒯 before sampling has occurred. That is ℱ, contains all of the leaves in 𝒯, plus additional extant and ancestral lineages that were not sampled. Following the reduced tree decomposition (e.g., Section 3.3 of Harris, Johnston, and Roberts 2020^95^), let us define a *stub* as an internal node in ℱ, conditional on one child lineage being sampled in 𝒯 and the other being unsampled (Fig. 2). Stubs appear in a time-dependent rate 2*λp*_0_(*t*) in 𝒯. Following Eqn (1) of Stolz, Stadler, and Vaughan 2024,^96^ the expected number of stubs along an interval [*t*_*o*_, *t*_*e*_] is:

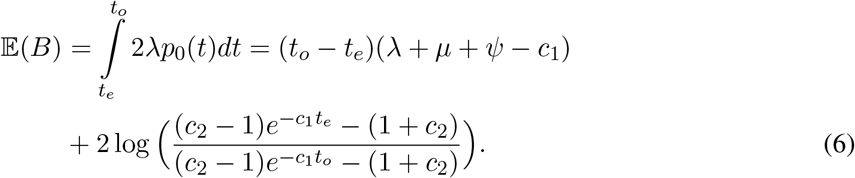

The number of stubs *B* follows a Poisson distribution with rate E(*B*). Given that stubs are conditional on one lineage being unsampled, they are more likely to occur further back in time (Fig. 3A). The probability density of observing *B* stubs at ages *Z* = (*z*_1_, *z*_2_, …, *z*_*B*_) along interval [*t*_*o*_, *t*_*e*_] is then:

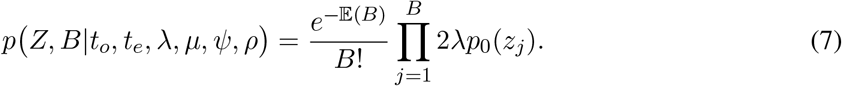

Lastly, let *B*_*i*_ and *Z*_*i*_ be the number and ages of stubs on branch *i*, and let 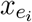 and 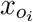 be the ages of branch *i* and its origin. Then, the overall probability density of observing a tree and its stubs is:

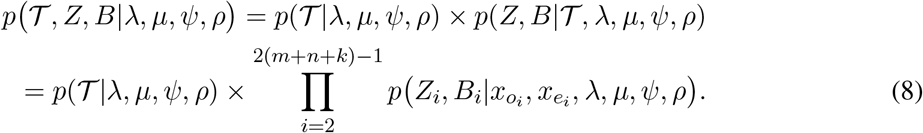

The product in the above equation starts at *i* = 2, since *i* = 1 corresponds to the root, and here we assume for simplicity that there are no stubs on the root branch (the origin). The number of stubs, their ages, and their branch indices are all estimated using reversible jump MCMC^97^ (Supporting Information).

### Clock models

Each internal branch *i* in 𝒯 is associated with a time duration *τ*_*i*_, a relative rate *r*_*i*_ and a spike *s*_*i*_ (Fig. 2). Spike sizes are independent of time, for example a spike of size 0.1 means that around 10% of sites in the alignment are expected to change per bifurcation, regardless of time units. The total evolutionary distance of a branch is then *µ*_*c*_ *× τ*_*i*_ *× r*_*i*_ + *s*_*i*_, where *µ*_*c*_ is the clock rate. These distances are used to compute the tree likelihood using Felsenstein’s peeling algorithm.^98^

Under the relaxed model of gradual evolution, each relative rate is independent and identically distributed with a mean of 1 so that it does not conflate with *µ*_*c*_:^4,43^

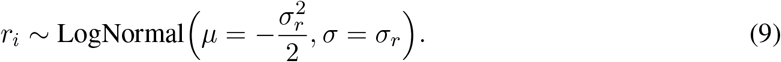

The standard deviation *σ*_*r*_ should be reasonably small (e.g., < 0.5) to ensure that these rates are constrained. This constraint is enforced by our prior distribution. We use the real-space rate parameterisation of this model (implemented in the BEAST 2 ORC package^43, 89^), which is built on top of the classic discretised form by Drummond et al. 2006.^4^

Under the relaxed model of abrupt evolution, the spike size of a branch is dependent on the number of stubs along that branch. This model has two hyperparameters: the spike mean *S*_*µ*_ and the spike shape *S*_*α*_. Each spike is independent and identically distributed under a gamma distribution with shape *k* and scale *θ*:

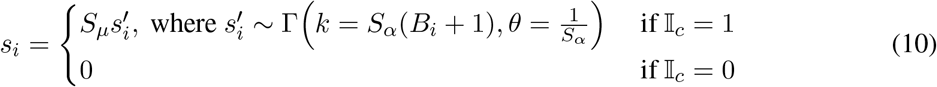

where *B*_*i*_ is the number of stubs on branch *i*, and 𝕀_*c*_ ∈ {0, 1} is the clock model indicator which determines whether we are using the gradual (𝕀_*c*_ = 0) or gradual+abrupt (𝕀_*c*_ = 1) model. This latter assumes that each bifurcation contributes an independent spike whose size is Gamma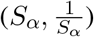distributed. The total branch spike sum is estimated rather than per-stub. Keeping the mean *S*_*µ*_ outside of the Gamma distribution (which has a mean of 1) helps to overcome numerical and mixing issues. Analogous to the standard deviation of the gradual relaxed clock, the shape *S*_*α*_ should be large (e.g., *>* 0.8), and therefore the variance small, so that spike sizes are constrained. Based on our estimates of spike size here, as well as previous ones,^25, 27, 39^ we recommend using a *S*_*µ*_ prior with a mean of around 0.01 in the general case where no information is known, e.g., LogNormal(mean=0.01, *σ* = 1.2).

Combining these clock models with the fossilised birth-death tree prior plus stubs (detailed in the previous section), the overall posterior density is:

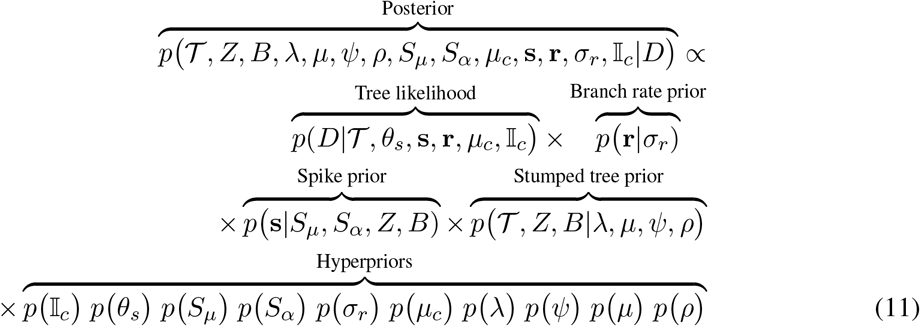

where *θ*_*s*_ contains parameters related to the site model (such as transition-transversion ratio in the case of DNA data, or amino acid frequencies in the case of protein). In all cases here, the clock model indicator has a 𝕀_*c*_ ∼Uniform({0, 1}) prior. As outlined in Supporting Information, we built upon an advanced pool of phylogenetic proposal kernels^43, 83, 86–89^ to ensure an efficient traversal of the posterior distribution, including tree topology, branch lengths, sampled ancestors, and stubs.

### Clade supports

The inner contours on the cocoa-fruit plots of Fig. 5 denote the statistically insignificant zones of clade support. This model assumes that the clade counts from either axis are independently drawn from a binomial distribution with *n* = 100 samples, and an unknown probability from a Uniform(0,1) prior.

The joint probability of observing 0 ≤ *k*_1_, *k*_2_ ≤ *n* observations is:

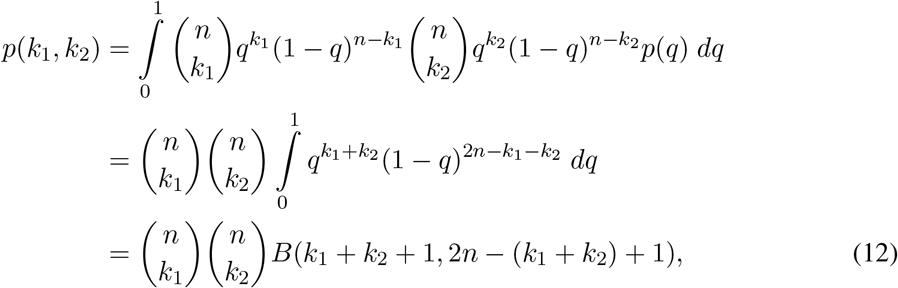

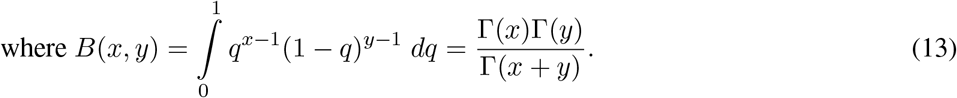

MCMC is run to achieve at least 200 (effective) samples, so 100 samples here is a conservative estimate, noting that more samples would give tighter bounds. In practice however, each clade support would have a distinct effective sample size (notably those with low support), and we currently have no robust means to quantify this. With these caveats noted, the inner contours of Fig. 5 are the 95% credible intervals of this distribution, and therefore represent statistically insignificant differences under this model.

The outer contours on Fig. 5 are the change in log Bayes factor. Following Kass and Raftery,^49^ a log_10_ Bayes factor between 0.5 and 1 is considered as substantial support of one hypothesis over another, 1-2 is strong, and over 2 is decisive The logarithm of the Bayes factor *b*_1_ for clade *C* under model *p*_1_ is:

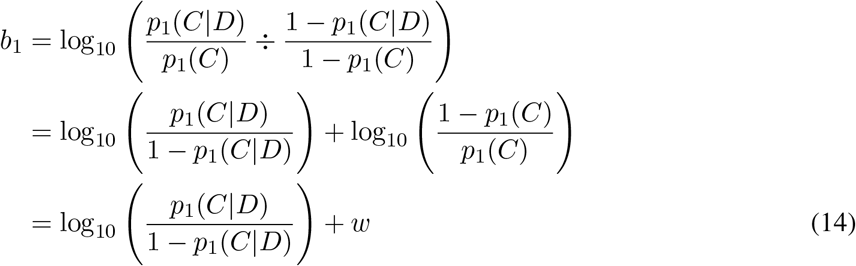

for sequence data *D*, where *p*_1_(*C*|*D*) and *p*_1_(*C*) are the posterior and prior support of *C* under model *p*_1_.

Now, suppose we have a second model *p*_2_ with the same tree prior distribution *p*_2_(*C*) = *p*_1_(*C*), but different likelihood *p*_2_(*C*|*D*)*/*= *p*_1_(*C*|*D*). Then, the change in log Bayes factor *H* is:

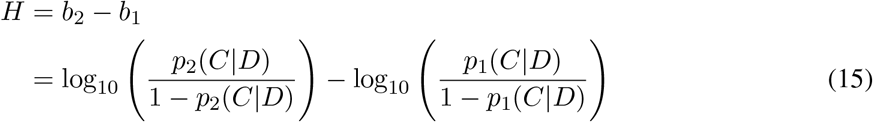

We can interpret *H* on the scale by Kass and Rafferty.^49^ For example, if the first model gives *p*_1_(*C*|*D*) = 0.5 and the second gives *p*_1_(*C*|*D*) = 0.59, then the log Bayes factor has increased by 0.16, which is a rather minor change. Now let us increase the probability by 0.09 again, but this time at an extreme end of the [0,1] domain. Suppose that *p*_3_(*C*|*D*) = 0.9 and *p*_4_(*C*|*D*) = 0.99 - in this case the latter model increases the log Bayes factor substantially, by 1.04; a 10^1.04^ = 11-fold change in Bayes factor. The outer contours on Fig. 5 depict the change in log Bayes factor, *H*, when subtracting the x-axis from the y-axis.

### Data, materials, and software availability

Our GPL-licensed source code is available as the `GammaSpikeModel` package for BEAST 2, where it is readily configured through a user-friendly interface. XML files, source code, and documentation are found at https://github.com/jordandouglas/GammaSpikeModel.

## Acknowledgements

This work was supported by the Alfred P. Sloan Foundation Matter-to-Life program Grant number G-2021-16944 and a New Zealand Royal Society Te Apārangi Marsden Fund (22-UOA-052). The au-thors gratefully thank Alexei Drummond for his helpful feedback on Bayesian inference and clade supports, and Padriac Amato Tahua O’Leary for providing explanatory insights behind the coupling between branching and evolution. The authors thank the artists of PhyloPic for creating the cichlid and cephalopod silhouettes (Fig. 1 and 4), which are available under public licenses at https://www.phylopic.org/.

