## Supporting Information for "Evolution is coupled with branching across many granularities of life"

September 9, 2024

### 1 MCMC proposal kernels

Proposal kernels, also known as operators, are the workhorses of MCMC. They are responsible for proposing a new state $\theta'$ conditional on the current state $\theta$. Putting extra time and thought into operator design can facilitate a faster traversal of the posterior distribution, which is often tricky to navigate in the case of large trees or long, signal-rich alignments (which may have "narrow" peaks in the likelihood landscape[1]). One of the greatest challenges in this task comes from ensuring that proposal kernels satisfy "detailed balance", which is a requirement for MCMC to correctly sample from the target distribution. The operators used by the gamma spike model have been built upon decades of research into this topic.

#### 1.1 Proposing new values for continuous parameters

We used the following standard BEAST 2 operators for proposing new values for real-space parameters, such as those related to branching, site models, or clock models:[2] `RandomWalk`, `Scale`, and `AVMN`. The former operators act on a single dimension of a single parameter, and are used in conjunction with the Bactrian proposal kernel for selecting jump size.[3] `AVMN` acts on several parameters simultaneously by learning their correlations.[4]

#### 1.2 Tree operators

Designing tree operators for this model required two additional considerations beyond standard BEAST 2 tree proposals. The first comes from sampled ancestors, which are non-extant taxa with branch length zero, representing an ancestor. We built off the tree operators designed by Gavryushkina et al. 2014,[5] which are not only aware of sampled ancestors, but also toggle non-extant taxa between being sampled ancestors or extinct. The second constraint stems from the dependency of stub heights on the tree. As detailed in Section 2, during reversible-jump MCMC, stub heights are estimated proportional to their branch length. Therefore, any change in tree topology or branch length may change the placement of a stub along its branch, and thus the Hastings ratio needs to account for this change. The gamma spike model package therefore uses special adaptations of the following tree operators, which are accounting for these two requirements. The second requirement only applies during reversible jump MCMC.

Standard BEAST 2 operators:

- `NarrowExchange`: moves a branch to a nearby position.
- `WideExchange`: moves a branch to a further position.
- `Uniform`: moves the height of a single internal node.

- TreeScale: moves the height of all nodes.
- RootScale: moves the height of the root.
- UpDown: moves the height of all nodes in the tree, as well as one or more continuous parameters that are correlated with the node heights.
- LeafToSampledAncestorJump: toggles a leaf between being a sampled ancestor (length 0) or not (length $> 0$).

We also extended the BICEPS operators, which are reported to be quite efficient on serially-sampled trees:[6]

- TreeFlex: scales all branch lengths within a confined epoch.
- TreeStretch: moves parts of the tree, while taking tip dates into account.

Lastly, the constant distance operator accounts for the correlation between branch rates and branch lengths, and has shown to greatly improve convergence in the relaxed clock model:[1,7]

- ConstantDistance: moves a node height, and then adjusts the rates on the tree branches around it such that the total genetic distance remains constant.

Together, these operators provided an effective traversal of tree space in a manner that ensures MCMC is correctly sampling the target distribution.

##### 1.3 Other spike model operators

- BitFlipOperator: toggles the clock model indicator $\mathbb{I}_c$ between 0 and 1.
- SpikeUpRateDown operator that first selects a branch, and then moves its rate up and its spike down, or vice versa. A Bactrian proposal kernel is used to select the extent of movement.[3] This is a special case of the UpDown operator.

##### 1.4 Stub operators

These are detailed in the following section.

#### 2 Estimating stubs

We describe three competing approaches to estimate stubs, each of which has been implemented in BEAST 2. Each method has unique advantages and disadvantages related to their their generalisability, computational efficiency, and in the extent of information made available during inference.

#### 2.1 Estimating the placement of each stub

The most complex and general method involves estimating the number of stubs $B$ on each branch index $\mathcal{I}$ as well as the ages of stub $Z$. To do this, we implemented a reversible-jump[8] proposal kernel for creating/destroying stubs in the tree. This proposal, termed `CreateOrDestroyStub`, is described in Algorithm 1. This implementation allows stubs to appear on any branch except for the root. To assist with convergence during MCMC, stub times are estimated relative to their position along their branch i.e., $0 < Z < 1$, and then transformed back into absolute times when computing the probability density function. The `Uniform` and `Scale` operators move existing stubs around their branch, by proposing a new value of $Z$, and the `Swap` operator exchanges two indices in the vector $\mathcal{I}$. These three proposals are standard operators in BEAST 2.[9]

We validated this method through coverage simulation studies.[10] Although it has the advantage of providing information on stub timing that may be useful for certain models, it comes at the cost of introducing further complexity during MCMC and hampering its convergence time.

#### 2.2 Estimating the number of stubs per branch

A faster approach involves estimating the number of stubs on each branch, and integrating over their times. Following Eqn (1) of Stolz, Stadler, and Vaughan 2024,[11] the expected number of stubs along an interval $[t_o, t_e]$ is:

$$\mathbb{E}(B) = \int_{t_e}^{t_o} 2\lambda p_0(t)dt = (t_o - t_e)(\lambda + \mu + \psi - c_1) + 2\log\left(\frac{(c_2-1)e^{-c_1 t_e} - (1+c_2)}{(c_2-1)e^{-c_1 t_o} - (1+c_2)}\right),$$

where all terms above are defined in Materials and Methods. The probability of observing $B$ stubs a branch with interval $[t_o, t_e]$ follows a Poisson distribution:

$$p(B|\delta) = \frac{\mathbb{E}(B)^B e^{-\mathbb{E}(B)}}{B!},$$

where $\delta = (\lambda, \mu, \rho, \psi)$ is the vector of branching parameters.

The operators used here are simple – a `RandomWalk` operator that randomly increments or decrements the number of stubs on a branch, and a `Swap` operator that swaps the number of stubs among two branches. This approach is general and allows the spike prior density to be computed (whose shape is proportional to $B + 1$). However, we have lost information on stub times.

#### 2.3 Stub-free inference (default setting)

The most efficient method involves integrating over stubs entirely (including the number of stubs and their placement in the tree). Let $S = S' \times S_\mu$ be the size of a spike, where $S'$ is the relative spike size from a Gamma(shape=$S_\alpha$, scale = $\frac{1}{S_\alpha}$) distribution with a mean of 1. The sum of spikes along a branch with $B$ stubs is thus:

$$p(S'|B,\delta) = p(S'|B) = \frac{1}{\Gamma(k)\theta^k}(S')^{k-1}e^{-S'/\theta}$$

where shape $k = S_\alpha(B+1)$ and scale $\theta = 1/(S_\alpha(B+1))$.

We do not know $B$ so we will integrate it out:

$$p(S'|\delta) \approx \sum_{B=0}^{N} p(S'|B,\delta)^{\mathbb{I}_c} \times p(B|\delta)$$

for some large $N$. We set $N$ such that the cumulative Poisson probability $\sum_B p(B|\delta)$ exceeds 0.999. If abrupt evolution is inactive ($\mathbb{I}_c = 0$), then we ignore the dependency on spikes. We note that when the number of stubs on a lineage is large, then the shape parameter $k$ becomes large, which can cause numerical issues.

We can now sample the number of stubs per lineage at the time of reporting estimates to the user (e.g., by logging every 10,000 MCMC samples). When logging the number of stubs $B$ on a lineage with relative spike size $S'$, inversion sampling is employed on the following distribution:

$$p(B|S',\delta) = \frac{p(S'|B,\delta)\ p(B|\delta)}{p(S'|\delta)}.$$

In contrast to the previous two approaches, there are no operators required to move stubs, because they are not part of the state. Although this method is the most efficient of the three for the spike model, it is not general. If the stubs were to be used in additional posterior distribution calculations then the above equation might not hold, and the reported number of stubs would be incorrect. Notwithstanding this potential fragility, this method remains the preferred one for the proposed model.

**Algorithm 1:** The `CreateOrDestroyStub` proposal kernel randomly creates or destroys a stub. This involves incrementing or decrementing the number of stubs $B$ and inserting/removing an entry from the vector of stub times $Z$ and the vector of branch indices $\mathcal{I}$. The proposal kernel has a Hastings-ratio that is dependent only on the time-length of the branch being operated on. Note that $n$ is the number of extant, $m$ extinct, and $k$ sampled ancestral taxa.

**Function** `CreateOrDestroyStub(`$B, Z, \mathcal{I}, \mathcal{T}$`)`**:**

- $A \sim \text{Uniform}(\{1, 2, \ldots, 2(n+m+k)-2\})$ ; `// Sample a non-root branch`
- $U \sim \text{Uniform}(0, 1)$ ; `// Create or destroy?`
- **if** $A$ is sampled ancestor **then**
  - **return** $B, Z, \mathcal{I}, 0$ ; `// Reject proposal`
- **if** $U < 0.5$ **then**
  - $B' \leftarrow B + 1$ ; `// Create stub on branch` $A$
  - $I \sim \text{Uniform}(\{0, 1, 2, \ldots, B\})$ ; `// Sample index to insert stub`
  - $R \sim \text{Uniform}(x_{e_A}, x_{o_A})$ ; `// Sample a stub time along` $A$
  - $Z' \leftarrow (z_1, z_2, \ldots, z_I, R, z_{I+1}, \ldots, z_B)$;
  - $\mathcal{I}' \leftarrow (\mathcal{I}_1, \mathcal{I}_2, \ldots, \mathcal{I}_I, A, \mathcal{I}_{I+1}, \ldots, \mathcal{I}_B)$;
  - HR $\leftarrow (x_{e_A} - x_{o_A})$ ; `// Hastings-ratio is the length of` $A$
- **else**
  - $F_A \leftarrow \sum_i \mathbb{I}(\mathcal{I}_i = A)$ ; `// How many stubs on` $A$`?`
  - **if** $F_A = 0$ **then**
    - **return** $B, Z, \mathcal{I}, 0$ ; `// Reject proposal`
  - $B' \leftarrow B - 1$ ; `// Destroy stub from branch` $A$
  - $I \sim \text{Uniform}(\{i | \mathcal{I}_i = A\})$ ; `// Sample eligible stub to destroy`
  - $Z' \leftarrow (z_1, z_2, \ldots, z_{I-1}, z_{I+1}, \ldots, z_B)$;
  - $\mathcal{I}' \leftarrow (\mathcal{I}_1, \mathcal{I}_2, \ldots, \mathcal{I}_{I-1}, \mathcal{I}_{I+1}, \ldots, \mathcal{I}_B)$;
  - HR $\leftarrow (x_{e_A} - x_{o_A})^{-1}$ ; `// Reverse Hastings-ratio`
- **return** $B', Z', \mathcal{I}'$, HR;

### 3 Direct simulation of fossilised-birth-death trees

We describe two methods for simulating a birth-death tree, and its stubs, directly (i.e., without inference under MCMC). In each case, the simulation starts with a single lineage and stops under varying criteria. After the simulation has ended, the $m$ extinct and $k$ ancestral lineages are sampled from the full phylogeny $\mathcal{F}$ by independently and randomly sampling points along the tree at rate $\psi$. These simulations assume that all extant taxa are sampled, i.e., $\rho = 1$.

#### 3.1 Standard approach

The standard approach assumes that one lineage branches into two at rate $\lambda$ and dies at rate $\mu$. The simulation is conditional on reaching $n$ extant lineages, and therefore any trees that become extinct before this point are discarded. A major limitation here is that tree simulation can be expensive and wasteful, especially under subcritical conditions $\mu > \lambda$ and when $n$ is large. Any internal nodes that are present in the full tree $\mathcal{F}$ but absent from the reduced/sampled tree $\mathcal{T}$ are then identified as stubs in a deterministic fashion, and their unsampled subtrees are discarded. This approach was used during our coverage simulation studies.

#### 3.2 Red-purple tree approach

We describe an alternate method that allows direct simulation of a birth-death tree conditional on the survival of at least one lineage at time $T$. This construct is restricted to sub- or super-critical conditions ($\lambda \neq \mu$) and when $\psi = 0$.

Let $N_s$ be the number of surviving lineages after $s$ units of time have elapsed after starting with one lineage. Then, the probability of all descendants being extinct is known to be:

$$q_s = \mathbb{P}(N_s = 0) = \frac{\mu\big(e^{(\lambda-\mu)s} - 1\big)}{\lambda e^{(\lambda-\mu)s} - \mu}. \tag{1}$$

Following the reduced tree decomposition (e.g., Harris, Johnston, and Roberts 2020[12]), let us colour a node born at time $t$ as purple if it has at least one descendent alive at time $T$, otherwise colour it red. A purple particle will split into two purple particles at time-dependent rate:

$$\lambda_b(t) = \lambda\ \mathbb{P}(\text{both offspring purple}|\text{parent purple}) = \lambda\frac{(1-q_{T-t})^2}{1-q_{T-t}} = \lambda(1-q_{T-t}). \tag{2}$$

This pure-birth process gives rise to the purple tree, also known as the reconstructed tree or the reduced tree. Purple particles can also give rise to red particles (this corresponds to an internal node where one child survives to time $T$ and the other does not). These nodes are what we call the stubs, and comprise the points where the red trees grow off the purple tree. A red node is born off a purple lineage at a time-dependent rate:

$$\begin{aligned}\lambda_r(t) &= \lambda\Big(\mathbb{P}(\text{left child red, right child purple}|\text{parent purple}) + \mathbb{P}(\text{left child purple, right child red}|\text{parent purple})\Big)\\ &= \lambda\,\frac{2q_{T-t}(1-q_{T-t})}{(1-q_{T-t})}\\ &= 2\lambda q_{T-t}. \end{aligned} \tag{3}$$

Similarly, red particles split into two red particles at rate $\lambda q_{T-t}$ and each red individual alive at time $t$ independently dies at rate $\mu/q_{T-t}$. The red-purple tree approach provides a method for directly simulating birth-death trees (including their stubs) that are guaranteed to survive, by starting with an initial purple particle, without the need for discarding trees that have gone extinct. This pure-birth process can be simulated by iteratively incrementing the time $t$ and then generating purple or red nodes at rates $\lambda_b(t)$ and $\lambda_r(t)$. In terms of computational efficiency, it is therefore particularly suitable for subcritical conditions $\mu > \lambda$.

### 4 Simulation studies

#### 4.1 Coverage simulation studies

We performed coverage simulation studies[10] to validate our phylogenetic model. This involved the following steps. First, we simulated a set of binary rooted time-trees $\mathcal{F}$ as birth-death processes, using $\lambda$, $\mu$, and $\psi$ sampled from their prior distributions (specified below). We used the ratio between $\lambda$ and $\mu$, i.e., the reproduction number, to ensure that $\lambda > \mu$, and $\rho$ was fixed at 1 during the simulation studies. These trees were simulated until the number of lineages reached $n$, as described above. Second, we pruned off all lineages that went extinct, as well as the origin branch, to arrive at the reduced tree $\mathcal{T}$. Third, we performed MCMC on each tree to estimate $\lambda$, $\mu$, $\psi$, and the stubs, using the same prior distribution that the parameters were initially sampled from. Trees were fixed to their true values in this step. Lastly, we compared the known values of $\lambda, \mu$, $\psi$, and $B$ with their posterior estimates. If this method is indeed well-calibrated, we would expect the true value to lie in the 95% credible interval approximately 95% of the time.

We validated our model with three coverage simulation studies. All three experiments provided confidence in the correctness of our method, and its ability to recover parameter estimates from data simulated under the same model used in inference.

**Experiment 1: Estimating tree from sampled branching parameters.** First, we compared the trees generated by our direct simulator with the distribution of trees estimated by MCMC (Fig. S1). If the simulator and inference machinery are sampling from the same probability distribution, then we would expect the simulated trees to be representative samples from the tree distribution inferred by MCMC, conditional on the branching parameters. 100 MCMC replicates were performed on 100 simulated trees, with 100 sets of independently sampled branching parameters. Each tree was simulated from a birth-death process, with stopping criteria of reaching $n = 20$ extant taxa (and a random number of extinct taxa $m$ and sampled ancestors $k$). Branching parameters were independently sampled from the following distributions during simulation, and then fixed to their true values during inference: $\lambda \sim \text{LogNormal}(\text{mean} = 2, \sigma = 0.5)$, reproduction number $\frac{\lambda}{\mu} - 1 \sim \text{Exponential}(\mu = 2)$, and sampling proportion $\frac{\psi}{\psi+\mu} \sim \text{Beta}(\alpha = 2, \beta = 5)$. The number and placement of stubs were also estimated.

This experiment provided confidence in the congruence between our two methods of sampling the tree probability distribution (simulation and and inference).

**Experiment 2: Estimating branching parameters from simulated trees.** Second, using these same 100 datasets, we estimated tree branching parameters from fixed trees that were directly simulated under those parameters (Fig. S2). The branching parameter distributions in the previous paragraph were used as prior distributions during inference here. This experiment, in conjunction with the one above, confirm the correctness of our method when the tree and/or branching parameters are known, and validate our new operator `CreateOrDestroyStub`.

**Experiment 3: Estimating branching parameters and tree from simulated DNA data.** Finally, we estimated all parameters – including trees, substitution model parameters, clock model parameters, and tree prior parameters – from nucleotide alignments simulated under the respective parameters (Fig. S3). The clock model was sampled during simulation, as gradual with probability 0.5, gradual+abrupt with probability 0.5. The clock model indicator was then estimated using the same probability distribution as the prior, allowing us to determine how often the true model can be recovered from the data. Clock model parameters were sampled from the following priors: gradual relaxed clock standard deviation $\sim$ Gamma$(\alpha = 5, \beta = 0.05)$, mean spike size $S_\mu \sim$ LogNormal$(\text{mean} = 0.05, \sigma = 0.8)$ substitutions per speciation, and spike gamma shape $S_\alpha \sim$ LogNormal$(\text{mean} = 2, \sigma = 0.5)$. These trees each contained $n = 40$ extant taxa and a variable number of sampled ancestral/extinct taxa, and were simulated from the following prior distributions: $\lambda \sim$ LogNormal$(\text{mean} = 10, \sigma = 0.5)$, reproduction number $\frac{\lambda}{\mu} - 1 \sim$ Exponential$(\mu = 2)$, and sampling proportion $\frac{\psi}{\psi+\mu} \sim$ Beta$(\alpha = 2, \beta = 10)$. Under this distribution, the mean tree height was 0.68 substitutions per site, with a 95% credible interval of (0.11, 1.9), while the mean total number of taxa (extant + extinct + sampled ancestral) was 48 (40, 67). Each nucleotide alignment was 100 sites and simulated under an HKY model with four gamma rate heterogeneity categories, with the following priors: transition-transversion ratio $\kappa \sim$ Exponential$(\mu = 5)$ and gamma rate heterogeneity shape $\sim$ Exponential$(\mu = 1)$. The four nucleotide frequencies were fixed at 0.25. 200 MCMC replicates were performed on 200 simulated datasets. This experiment provide further confidence in our method and also demonstrate that even just short sequences (100 nucleotides) contain enough information to confidently identify the true clock model using our MCMC Bayesian model averaging procedure. The true values were generally well-correlated with the simulated values (Pearson's correlation coefficient ranging from 0.5 to 0.99), and would likely become more correlated with larger datasets.

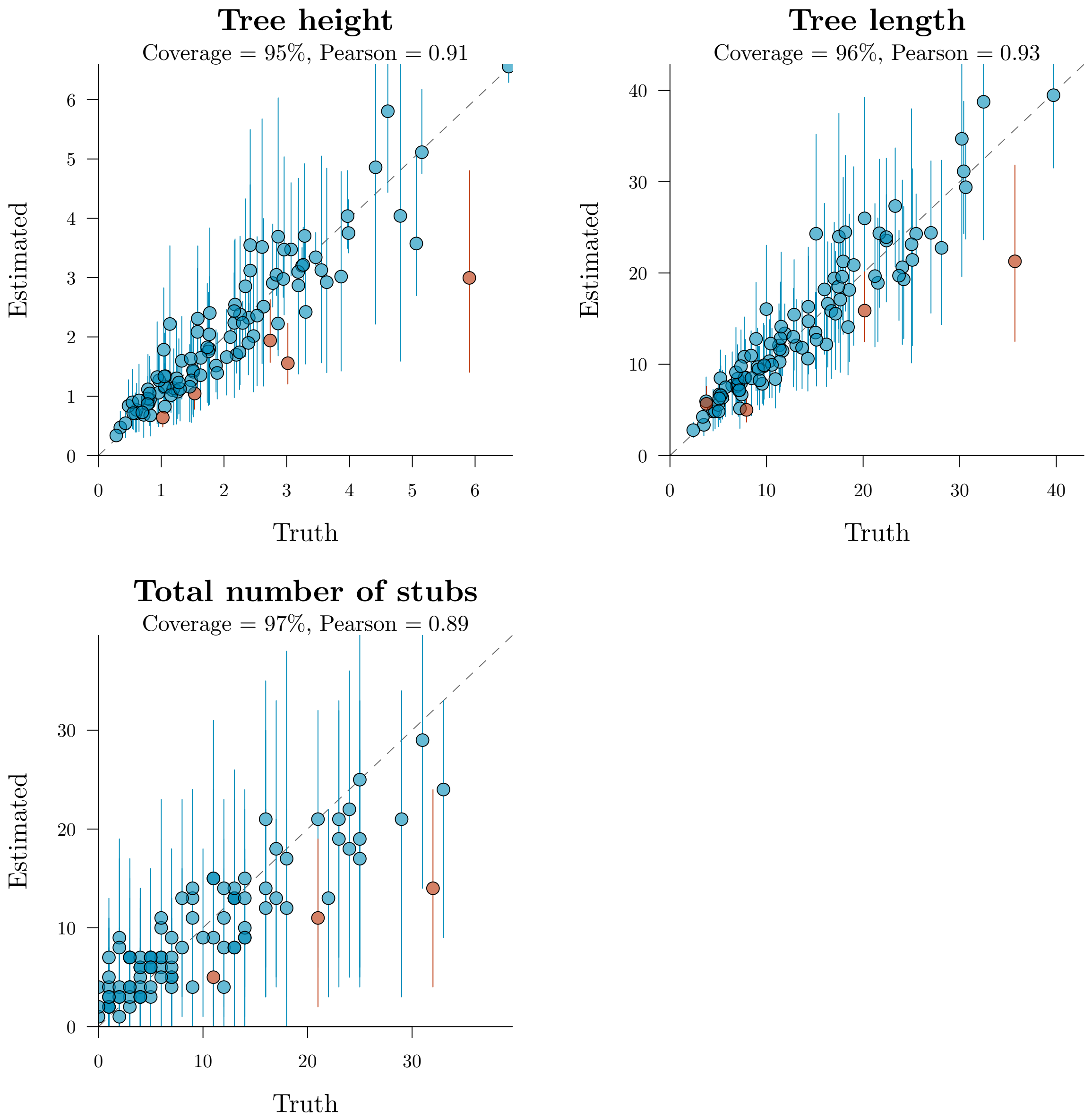

Fig. S1: Experiment 1 – Estimating trees from known branching parameters. Each point is an independent MCMC replicate on an independently simulated tree, including its branching parameters (which are fixed). Circles represent mean estimates under the posterior distribution, and vertical lines are 95% credible intervals. A coverage close to 95% means that around 95% of true values are contained within their 95% credible interval, thus providing confidence in correctness of the method. The Pearson correlation quantifies the agreement between the true values and their mean estimates.

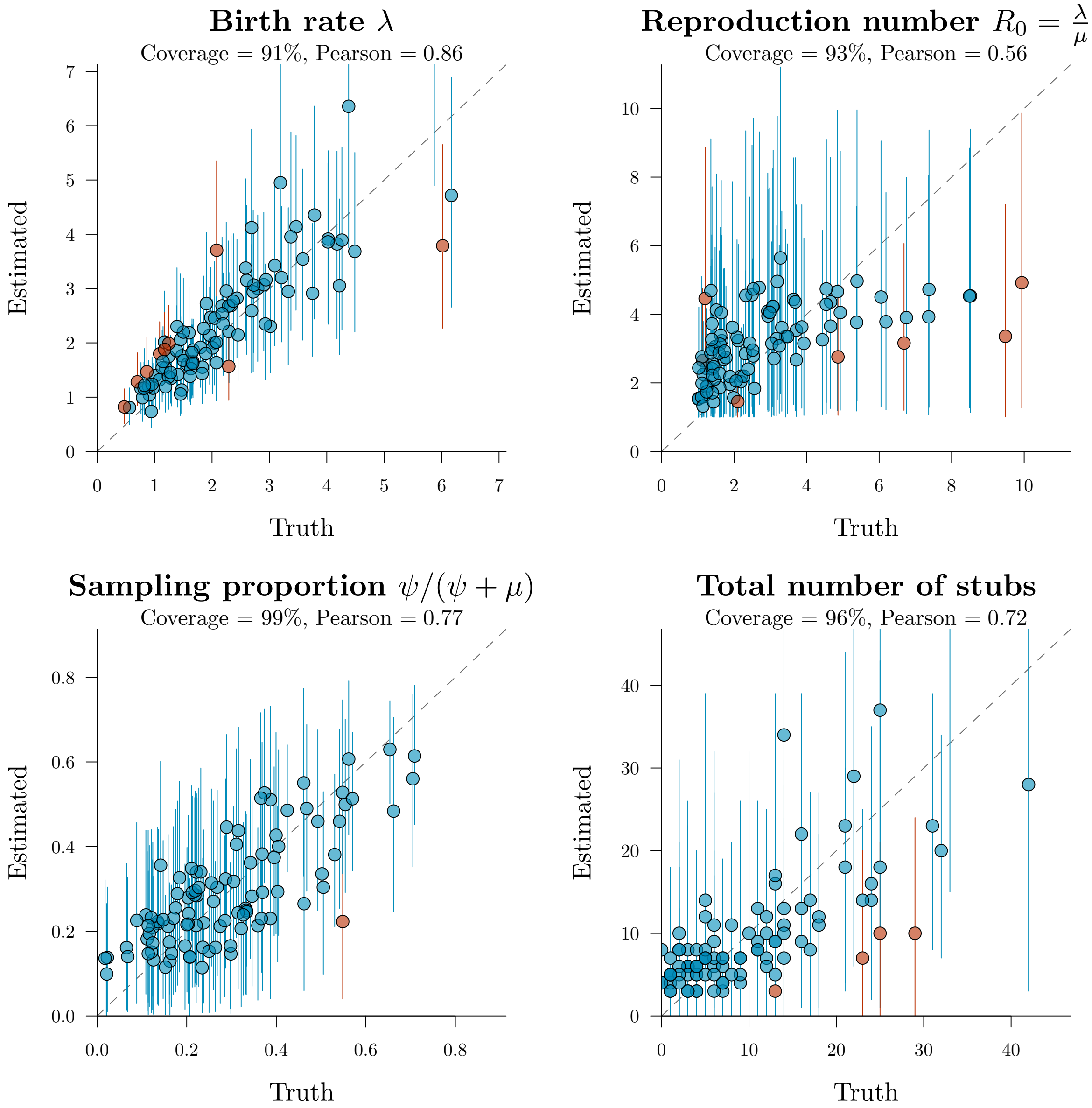

Fig. S2: Experiment 2 – Estimating branching parameters from known trees. In contrast to Fig. S1, the trees here are fixed at their true topologies and lengths, while branching parameters and stubs are estimated.

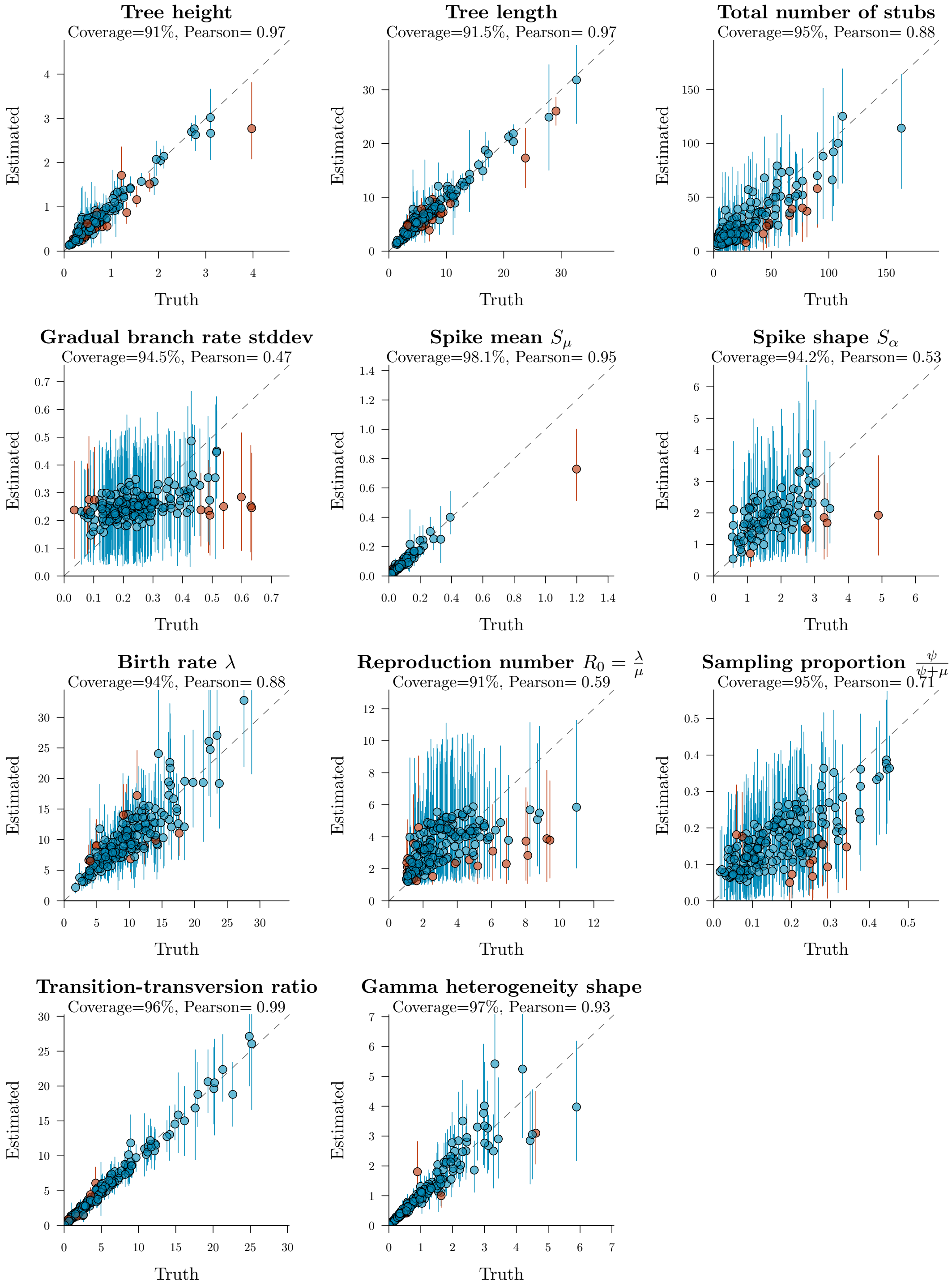

Fig. S3: Experiment 3 – Validating the two clock models. Each point is an MCMC replicate on an independently simulated nucleotide dataset (see Fig. S1). The tree height and length are the root height and branch length sum, respectively (units of substitutions per site). The spike mean and shape posteriors are conditional on the gradual+abrupt model being estimated, and hence their plots have fewer points.

#### 4.2 Bayesian model averaging

By performing additional simulation studies, we characterised how accurately our new method can identify the true clock model (gradual or gradual+abrupt) on data simulated under a known clock model. To do this, we simulated datasets under a random clock model (gradual with 0.5 probability, and abrupt with 0.5 probability) and estimated the clock model indicator $\mathbb{I}_c$ during MCMC. The simulations were conducted under the same conditions described in **Experiment 3**, with at least 40 taxa per tree, but with $S_\mu$ fixed at 0.01 during simulation and estimated during inference under a LogNormal(mean=0.01, $\sigma = 0.8$) prior. As shown in Fig. S4 (and Fig 3D of the main article), our new method can often recover the true clock model with high confidence, even on small datasets. The method is sometimes uncertain but very rarely wrong.

To test if the method is statistically consistent, we assessed simulated nucleotide alignments of varying lengths (20, 100, and 500 sites). These results confirm that increasing the amount of data also increases the chances of identifying the correct clock model (Fig. S4). Thus, the method appears to be statistically consistent. 20 sites is clearly too few for the method to be useful when the spike size is 0.01, but even under these conditions, the model is still rarely wrong.

To confirm that the signal for identifying the correct model came from the data and not from the prior distributions, we changed the prior distributions used during inference, as described below. These changes were quite extreme and yielded uninformative priors. They resulted in a greater amount of uncertainty in identifying the true clock model (Fig. S4). However, this uncertainty diminished with increasing amounts of data. Moreover, the method is very rarely wrong even with uninformative (and mismatching) prior distributions. This confirms that even short alignments still carry enough information to discriminate between different modes of evolution, even with little *a priori* knowledge.

- Birth rate $\lambda$:
  - $\text{LogNormal}(\text{mean} = 10, \sigma = 0.5)$ during simulation
  - $\text{LogNormal}(\text{mean} = 10, \sigma = 3)$ during MCMC (uninformed prior)
- Reproduction number $\frac{\lambda}{\mu} - 1$:
  - $\text{Exponential}(\mu = 2)$ during simulation
  - $\text{Exponential}(\mu = 5)$ during MCMC (greater mean and variance)
- Sampling proportion $\frac{\psi}{\psi+\mu} - 1$:
  - $\text{Beta}(\alpha = 2, \beta = 10)$ during simulation
  - $\text{Beta}(\alpha = 2, \beta = 2)$ during MCMC (uninformed prior)
- Mean spike size $S_\mu$:
  - $0.01$ during simulation
  - $\text{LogNormal}(\text{mean} = 0.1, \sigma = 2)$ during MCMC (uninformed prior)
- Spike shape $S_\alpha$:
  - $\text{LogNormal}(\text{mean} = 2, \sigma = 0.5)$ during simulation
  - $\text{LogNormal}(\text{mean} = 1, \sigma = 2)$ during MCMC (uninformed prior)

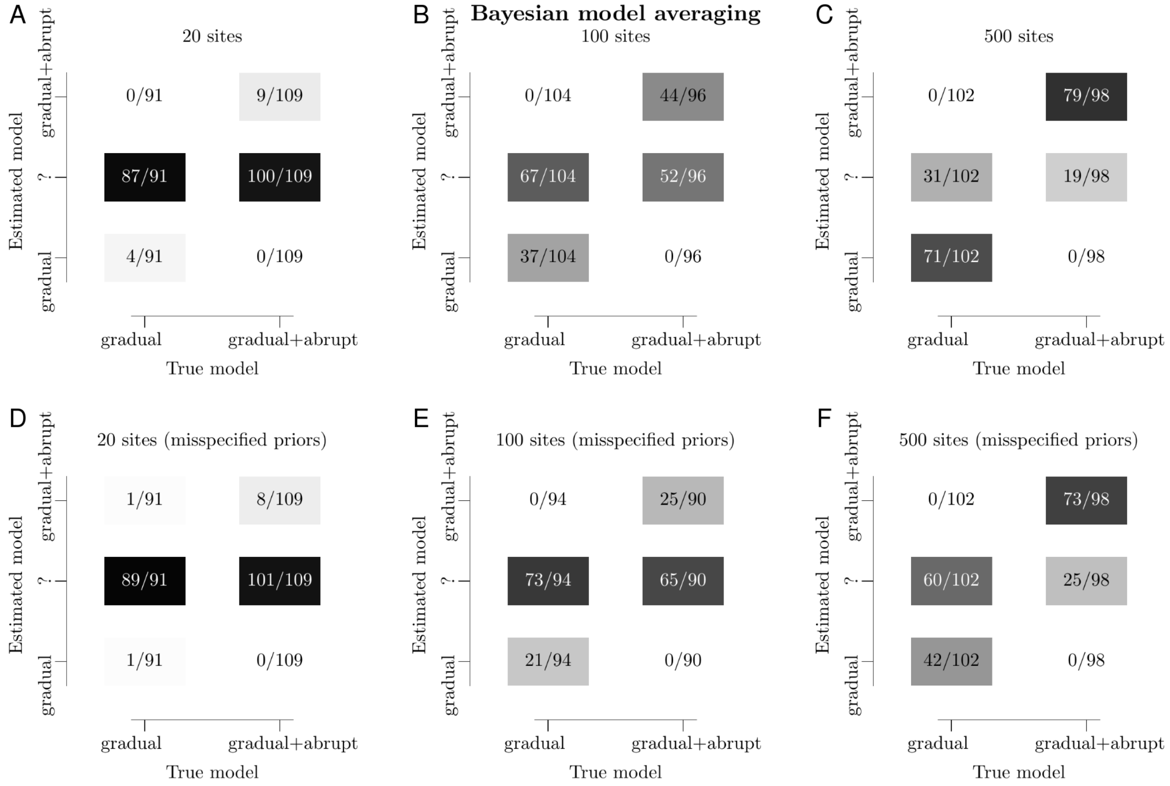

Fig. S4: Assessing how clock model estimation responds to varying alignment sizes and model misspecification. In each panel, 200 datasets were simulated under a clock model (x-axis) and then classified into a clock model using MCMC (y-axis). The '?' denotes the uncertain case where a clock model cannot be identified with a Bayes factor over 10 (i.e., $0.09 < p(\mathbb{I}_c = 1) < 0.91$). On the top row (A, B, and C), the same prior distributions were used during simulation and inference. The bottom row (D, E, and F) used the same datasets as above, however inference was done under a very different set of prior distributions, reflecting a more realistic scenario where the user does not have a solid grasp on *a priori* information (i.e., model misspecification). These experiments confirm that even short multiple sequence alignments carry rich information about whether they were generated through abrupt evolution.

##### 4.3 Assessing the effect of model misspecification on estimated tree length

As presented in Fig. 3E and 3F of the main article, we also assessed the effect of doing inference under the wrong clock model. This work involved simulating datasets under one model (with spikes) and then doing inference under a second model (with or without spikes). This experiment detected biased estimated of tree heights when the gradual model is used on spiked data. The most important parameter here is the spike mean $S_\mu$ – if this parameter was lower then the bias would be smaller.

- Clock rate $\mu_C$:
  - Fixed at 1 during simulation
  - LogNormal$(\text{mean} = 1, \sigma = 0.5)$ during MCMC (informed prior)
- Mean spike size $S_\mu$:
  - LogNormal$(\text{mean} = 0.02, \sigma = 0.8)$ during simulation
  - LogNormal$(\text{mean} = 0.05, \sigma = 0.8)$ during MCMC (slightly biased prior)
- Birth rate $\lambda$:
  - LogNormal$(\text{mean} = 5, \sigma = 0.5)$ during simulation
  - LogNormal$(\text{mean} = 1, \sigma = 3)$ during MCMC (uninformed prior)
- Reproduction number $\frac{\lambda}{\mu} - 1$:
  - Exponential$(\mu = 3)$ during simulation
  - Exponential$(\mu = 2)$ during MCMC (slightly biased prior)
- Sampling proportion $\frac{\psi}{\psi+\mu} - 1$:
  - Beta$(\alpha = 10, \beta = 4)$ during simulation
  - Beta$(\alpha = 2, \beta = 2)$ during MCMC (uninformed prior)
- Spike shape $S_\alpha$:
  - LogNormal$(\text{mean} = 2, \sigma = 0.5)$ during simulation and MCMC
- Gamma rate heterogeneity shape
  - Exponential$(\mu = 1)$ during simulation and MCMC
- Transition-transversion ratio
  - Exponential$(\mu = 5)$ during simulation and MCMC

Under these conditions, the trees each had $n = 20$ taxa and a random number of extinct and sampled-ancestral taxa: a mean of 40.3 and a 95% credible interval of [1, 121]. Their heights were 0.97 (0.19, 2.4) gradual substitutions per site. Nucleotides were simulated under an HKY model with four gamma rate heterogeneity categories.

### 5 Empirical datasets

#### 5.1 Aminoacyl-tRNA Synthetases

The following prior distributions were used for the aaRS analyses:

- Birth rate $\lambda \sim \text{LogNormal}(\text{mean} = 30, \sigma = 1)$
- Reproduction number $\frac{\lambda}{\mu} - 1 \sim \text{Exponential}(\mu = 5)$
- Sampling rate $\psi = 0$ (all taxa are extant)
- Clock rate $\mu_c = 1$ (all taxa are extant)
- Gradual relaxed clock standard deviation $\sim \text{Gamma}(\alpha = 5, \beta = 0.05)$
- Spike mean $S_\mu \sim \text{LogNormal}(\text{mean} = 0.01, \sigma = 1.2)$
- Spike shape $S_\alpha \sim \text{LogNormal}(\text{mean} = 2, \sigma = 0.5)$
- Gamma rate heterogeneity shape $\sim \text{Exponential}(\mu = 1)$

We used the WAG amino acid substitution model[13] with four gamma rate categories.

#### 5.2 Cephalopoda

The following prior distributions were used for the Cephalopod analyses:

- Birth rate $\lambda \sim \text{LogNormal}(\text{mean} = 10^{-3}, \sigma = 2)$
- Reproduction number $\frac{\lambda}{\mu} - 1 \sim \text{LogNormal}(\text{mean} = 0.2, \sigma = 1)$
- Sampling rate $\psi \sim \text{Beta}(\alpha = 1, \beta = 10)$
- Clock rate $\mu_c \sim \text{LogNormal}(\text{mean} = 10^{-4}, \sigma = 2)$ substitutions per site per million years
- Gradual relaxed clock standard deviation $\sim \text{Gamma}(\alpha = 5, \beta = 0.05)$
- Spike mean $S_\mu \sim \text{LogNormal}(\text{mean} = 0.01, \sigma = 1.2)$
- Spike shape $S_\alpha \sim \text{LogNormal}(\text{mean} = 2, \sigma = 0.5)$
- Gamma rate heterogeneity shape $\sim \text{Exponential}(\mu = 1)$

Alignment sites were partitioned into five Mk substitution models[14] based on the number of states (2, 3, 4, 5, and 6). The relative evolutionary rate of each site model was estimated, weighted by the number of sites in that site model. Each site model had four gamma rate heterogeneity categories. Fossil tip dates were used as described by Whalen and Landman 2022.[15]

#### 5.3 Indo-European languages

The following prior distributions were used for the Indo-European analyses:

- Birth rate $\lambda \sim \text{LogNormal}(\text{mean} = 0.7, \sigma = 0.1)$
- Reproduction number $\frac{\lambda}{\mu} - 1 \sim \text{Exponential}(\mu = 0.2)$
- Sampling rate $\psi \sim \text{Beta}(\alpha = 2, \beta = 100)$
- Clock rate $\mu_c \sim \text{LogNormal}(\text{mean} = 10^{-3}, \sigma = 2)$ substitutions per site per thousand years
- Gradual relaxed clock standard deviation $\sim \text{Gamma}(\alpha = 5, \beta = 0.05)$
- Spike mean $S_\mu \sim \text{LogNormal}(\text{mean} = 0.01, \sigma = 1.2)$
- Spike shape $S_\alpha \sim \text{LogNormal}(\text{mean} = 2, \sigma = 0.5)$
- Covarion alpha $\sim \text{Exponential}(\mu = 1)$
- Covarion switch rate $\sim \text{Exponential}(\mu = 1)$
- Gamma rate heterogeneity shape $\sim \text{Exponential}(\mu = 1)$

Cognates were assumed to evolve under a binary covarion model[16] with four gamma rate categories and uniform hidden frequencies (0.5, 0.5). Ancient language dates were estimated based on informed prior distributions, as described by Heggarty et al. 2023.[17]

#### 5.4 Benchmark datasets

We tested our method to the nucleotide datasets DS1 – 11 by Whidden and Matsen 2015.[18] These datasets are frequently applied for testing phylogenetic methods. As shown in Table S1, the spike model was favoured (with over 90%) support, on five datasets, and rejected (with under 10% support) on three. There was an intermediate level of support for the remaining three. When the spike model was favoured, the mean spike size was quite variable: ranging from 0.005 (DS6) to 0.02 (DS2). These estimates are quire similar to those made on our three focal datasets in the main article. Unsurprisingly, the datasets that rejected abrupt evolution had quite small estimates of $S_\mu$ when they were estimated. All chains were run until the effective sample sizes exceeded 200. For DS6 and DS10, this process was facilitated by coupled MCMC.[19] Much like the three case studies in the main article, the eleven datasets here each displayed a small (but negative) correlation between the estimated spike and rate of each branch.

| Data | Ntaxa | Nsites | Study | $p(\mathbb{I}_c = 1)$ | Spike mean $S_\mu$ | Stubs | $\rho_{sr}$ |
|---|---|---|---|---|---|---|---|
| **DS1** | 27 | 1949 | Hedges (1990)[20] | 1.0 | 0.0061 (0.0028, 0.0097) | 5.3 [0, 18] | -0.13 |
| **DS2** | 29 | 2520 | Garey (2012)[21] | 0.92 | 0.023 (0.00011, 0.084) | 4.6 [0, 13] | -0.049 |
| DS3 | 36 | 1812 | Yang and Yoder (2003)[22] | 0.67 | 0.0052 (0.00028, 0.013) | 8.3 [0, 22] | -0.056 |
| **DS4** | 41 | 1137 | Henk (2003)[23] | 0.93 | 0.0096 (0.00055, 0.021) | 10 [0, 29] | -0.13 |
| DS5 | 50 | 378 | Lakner (2008)[24] | 0.83 | 0.019 (0.00044, 0.033) | 18 [0, 120] | -0.1 |
| **DS6** | 50 | 1133 | Zhang and Blackwell (2001)[25] | 0.99 | 0.0046 (0.0026, 0.0069) | 29 [1, 65] | -0.066 |
| DS7 | 59 | 1824 | Yoder and Yang (2004)[26] | 0.0025 | 0.0038 (2.8e-05, 0.00052) | 170 [24, 320] | -0.038 |
| DS8 | 64 | 1008 | Rossman (2001)[27] | 0.65 | 0.0021 (0.00018, 0.0042) | 16 [0, 45] | -0.063 |
| DS9 | 67 | 955 | Ingram and Doyle (2004)[28] | 0.0013 | 2e-04 (2.4e-05, 0.00036) | 26 [1, 70] | -0.019 |
| **DS10** | 67 | 1098 | Suh and Blackwell (1999)[29] | 0.95 | 0.0058 (0.00074, 0.0096) | 5.6 [0, 17] | -0.091 |
| DS11 | 71 | 1082 | Kroken and Taylor (2000)[30] | 0.00025 | 0.00019 (2.8e-05, 0.00033) | 480 [200, 770] | -0.077 |

Table S1: Running the gradual-abrupt model on test datasets.[18] The spike model probability $p(\mathbb{I}_c = 1)$ was estimated using Bayesian model averaging, while $S_\mu$ and the number of stubs were estimated under the gradual+abrupt model, i.e., $\mathbb{I}_c = 1$. Datasets are in bold where abrupt evolution has over 90% support. Mean estimates (and 95% credible intervals) are shown. $\rho_{sr}$ is the average Pearson correlation between the estimated spike and rate of each branch, conditional on $\mathbb{I}_c = 1$. Note the differences between this table and Table 1 in the main article: i) here $p(\mathbb{I}_c = 1)$ is explicitly reported, while the datasets in Table 1 have $p(\mathbb{I}_c = 1) = 1$ in all three cases, and ii) here we do not report the results of the gradual model (only gradual+abrupt).
